# LIN28-mediated gene regulatory loops synchronize developmental transitions throughout organogenesis

**DOI:** 10.1101/2025.01.22.634324

**Authors:** Indhujah Thevarajan, Maria F. Osuna, Sonia Fuentes Lewey, Eustolia Sauceda, Sayra Briseno, Caylah Griffin, Bareun Kim, R. Grant Rowe, Edroaldo Lummertz da Rocha, Jihan K. Osborne

**Affiliations:** Department of Pharmacology, University of Texas Southwestern Medical Center, 5323 Harry Hines Blvd., Dallas, TX 75390, USA; Division of Pediatric Hematology/Oncology, Boston Children’s Hospital Boston, MA 0211; Department of Microbiology, Immunology and Parasitology (MIP), Centro de Ciências Biológicas, Universidade Federal de Santa Catarina Florianópolis, SC – Brazil; Simmons Comprehensive Cancer Center University of Texas Southwestern Medical Center, 5323 Harry Hines Blvd., Dallas, TX 75390, USA

## Abstract

Precise control of the intervals between self-renewal, proliferation, and differentiation of stem/progenitor cells are coordinated by developmental regulators, comprised of both microRNAs (miRNAs) and proteins, termed heterochronic genes. These heterochronic factors make up a unique subset of evolutionarily conserved genes that regulate the developmental rate and timing of metazoans from worms to mammals. We and others have shown critical roles for the RNA-binding proteins (RBPs) Lin28 during pluripotency, reprogramming, and organogenesis. There has been much investigation into the negative feedback loop between the Lin28-RBPs and the miRNAs–*Let-7* during development and disease. Albeit there are fewer investigations into how positive feedback loops between mammalian Lin28-RBPs and mRNAs order mammalian spatiotemporal transitions of progenitors from specification to organogenesis. Screening for factors that activate luciferase reporters of the human *LIN28A* and *LIN28B* promoters, in combination with genetic mouse models, we demonstrate positive feedforward loops between key developmental transcription factors such as B-Catenin, Sox2, Sox9, and Lin28-RBPs. Furthermore, we demonstrate heterochronic regulation of morphogenesis and ultimately differentiation is not only genetically moderated but also molecularly fine-tuned via position-dependent sequences in the 5’ and/or 3’ untranslated regions.

## Introduction

Genetic changes that modulate the developmental timing of an organism–1) in relation to other organisms in the same genus, and/or –2) with regards to the cell fate of one tissue relative to other tissues in the same organism are classified as heterochrony (Ambros and Horvitz 1984; McNamara 2012). The rate and timing of development is regulated by a subset of factors identified as heterochronic genes (Ambros and Horvitz 1984). Originally discovered in *C.* e*legans* as one of the key axes that orchestrates the intervals of proliferation and the transition to differentiation (Ambros, 1997), these gene regulatory networks consisted of the first miRNAs discovered – *lin-4* followed by *let-7;* and DNA/RNA-binding proteins – *hbl-1*, *lin-14* and *lin-28* (Ambros and Horvitz 1984; Lee et al. 1993). The temporally regulated feedback loop between the RNA-binding protein (RBP) Lin28 and the miRNA *let-7* is evolutionarily conserved in all metazoans. Loss of both Lin28 paralogs leads to early embryonic lethality in mice (Shinoda et al. 2013b), and human GWAS studies associates SNP variations of LIN28 with the improper timing of puberty (Zhu et al. 2010; Corre et al. 2016). In the early embryo, Lin28 binds to the stem-loop of the *let-7’s,* obstructing their processing and subsequent expression. At later stages of development, once expressed, *let-7* binds the 3′ untranslated regions (UTRs) of the *Lin28* mRNAs to suppress their translation (Piskounova et al. 2008; Rybak et al. 2008; Viswanathan et al. 2008). Studies in worms showed that Lin-28 functions via two genetically distinct mechanisms. The first mechanism, via Lin-28 binding to the 3’UTR of *hbl* mRNA, leading to its enhanced translation at the L1-L2 transition, with the second mechanism occurring at the L2-L3 transition via suppression of the *Let-7* miRNAs (Vadla et al. 2012).

The Lin28 paralogs (Lin28A and Lin28B) were previously shown to be strict onco-fetal proteins whose expression, via a reciprocal regulatory relationship with the family of *Let-7* miRNAs, are high throughout early development, off in adult tissues, and re-expressed in various tumors. Unlike many oncogenes, the amplification and mutation rates of LIN28A/B are extremely low, which is suggestive of a transcriptional re-activation (Viswanathan et al. 2009). We and others have previously demonstrated expression of Lin28a and Lin28b throughout the whole murine embryo that persists in the cell lineages of all three germ layers until about E9.5, after which is restricted to certain tissues, such as the lung (Yang and Moss 2003; Osborne et al. 2021). We have recently demonstrated that these mammalian paralogs are also heterochronic regulators of branching morphogenesis in the lung, kidney, and mammary glands, and that when knocked out in the lung epithelium, lead to perinatal lethality (Yermalovich et al. 2019; Osborne et al. 2021). Similar to the two distinct RNA regulatory mechanisms found in worms, we observed the actions of Lin28a and Lin28b in the early embryonic lung were *Let-7*-independent in regulating the mRNA targets *Etv5, Sox2,* and importantly *Sox9* (Osborne et al. 2021).

Lung development begins with the patterning and differentiation of the definitive endoderm into the anterior foregut and posterior midgut organ domains (Wells and Melton 1999; Serls et al. 2005). The forkhead transcription factors, specifically Foxa2 and Foxa3, regulate the induction of foregut and midgut endodermal patterning (Ang et al. 1993; Monaghan et al. 1993; Burtscher and Lickert 2009). Once the lung is specified, the first transcription factor that is expressed and regulates proper lung development is Nkx2-1 (Kimura et al. 1996; Minoo et al. 1999). The developed lung epithelium becomes functionally divided into two divergent compartments: the bronchiolar conducting airways of the proximal lung and the alveolar gas exchange region of the distal lung. Sox2 and Sox9 have been shown to mark and control the two developmental waves of lung branching morphogenesis (Gontan et al. 2008; Tompkins et al. 2009; Chang et al. 2013; Alanis et al. 2014). The first wave produces branches that develop into both proximal and distal lineages and is marked by Sox9, while the second wave regulates specification of the proximal region, demarcating the bronchoalveolar junctions, and is marked by Sox2 (Alanis et al. 2014). Correct lung formation is contingent upon reciprocating paracrine and autocrine signaling of the endodermal-derived epithelium and the mesodermal-derived mesenchyme (Herriges and Morrisey 2014). Several key signaling pathways together with multiple downstream transcriptional effectors, such as Wnt/B-Catenin and Fgf10/Etv5, Etv4, and Sox9, were found to cooperate in feedback and feedforward regulation, one example being B-Catenin upregulation of *Sox9* expression during lung specification (Ostrin et al. 2018).

In our present study, we conducted a luciferase reporter screen for transcription factors that were able to transactivate the human *LIN28A/LIN28B cis-*regulatory elements. We found the pluripotency factors OCT4, SOX2, KLF4, NANOG, MYC, along with factors that control multipotent progenitors (such as MYCN and B-CATENIN), hematopoiesis (such as MYB), endoderm differentiation (regulated by FOXA2 and FOXA3), mesoderm/neuromesodermal differentiation (via NOTCH and HOXB4), and epithelial lung development (via NKX2-1 and SOX9) were all able to differentially increase expression and the luciferase activity of *LIN28A* and *LIN28B.* Previously, we demonstrated the ability of LIN28A protein to bind and regulate the stability and the translation of *Sox9, Etv5,* and to a lesser extent *Sox2* mRNA (Osborne et al. 2021). In our current study, we found several mRNAs of transcription factors able to regulate the human *LIN28A/LIN28B* enhancer/promoter elements in our reporter screen were also present in the polyribosome fractionation of brain-specific double knockout (dKO) and differentially expressed in lung-specific dKO of Lin28a/b. Finally, we observed lung epithelial-specific knockout of B-Catenin, Sox2, and Sox9 at the onset of branching morphogenesis led to a significant decrease of *Lin28a,* but not *Lin28b,* mRNA expression.

## Results

### -2kb from TSS of the *Lin28A* and *Lin28B* genomic regulatory sequences more active than other putative sequences

Phylogenic analysis suggests a gene duplication event in the ancestral invertebrate heterochronic gene, *lin-28* (originally found in C. *Elegans*), led to the appearance of the vertebrate paralogs, Lin28a and Lin28b (Guo et al. 2006; Ouchi et al. 2014). Previous protein sequence alignments of *Drosophila, Xenopus,* Zebrafish, and mammals confirmed the presence of two unique RNA-binding domains: a cold shock domain (CSD), and two CCHC zinc-finger motifs (not usually found together in animals) however, found in the plant Lin28 homologue, PpCSP1 (Moss and Tang 2003; Ouchi et al. 2014; Li et al. 2017). Vertebrate Lin28A proteins clustered together with strong bootstrap support, while Lin28B formed a separate clade that is slightly more divergent (Fig.1A). Interestingly, early branching vertebrates – such as lamprey– appear to still have a single Lin28 that did not cluster with Lin28A nor was completely Lin28B-like. While protein sequences imply one level of conservation, the genomic regulatory sequences suggest another. Lin28A/B expression is important for the maternal oocyte-zygote transitions (West et al. 2009; Vogt et al. 2012; Sun et al. 2022), germ layer specification/early embryonic development (West et al. 2009; Cai et al. 2013; Faas et al. 2013; Shinoda et al. 2013a; Shinoda et al. 2013b; Zhang et al. 2016; Gundermann et al. 2019), neuroectoderm differentiation, and more recently, kidney and lung branching morphogenesis (Yermalovich et al. 2019; Osborne et al. 2021).

**Fig. 1.**
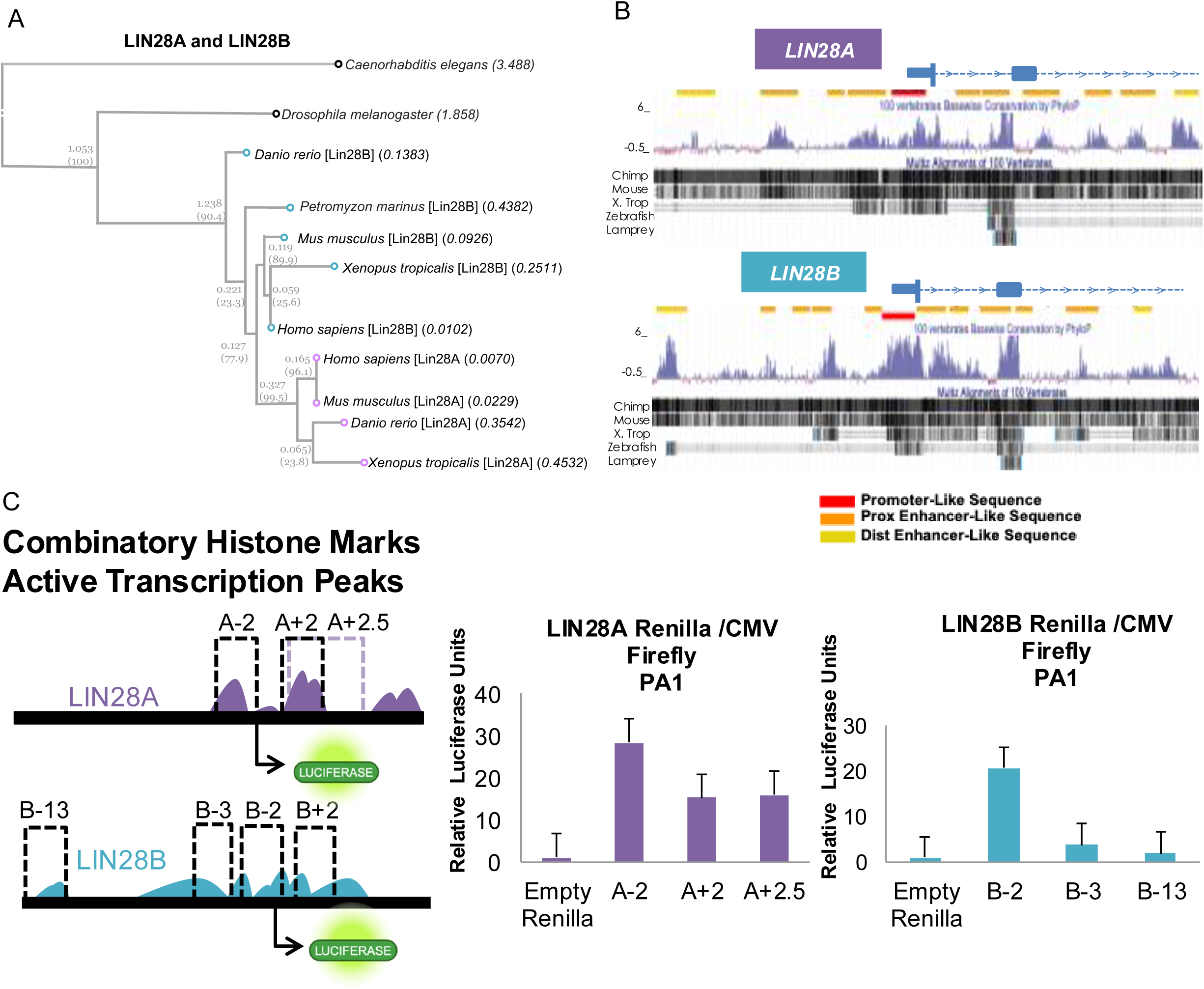
Comparison of protein and genomic regulatory sequences. A) Phylogenic analysis of Lin28A and Lin28B protein sequences using the bioinformatics tool https://www.genome.jp/en/gn_tools.html >Tree>phylogeny. B) Visualization of conserved genomic regulatory sequences for human *LIN28A* and *LIN28B,* in Chimp, Mouse, *Xenopus,* Zebrafish and Lamprey using UCSC genome browser (Perez et al. 2024). Assembly used was hg38 for *LIN28A* (chr1:26,406,455-26,417,607) RefSeq NM_024674.6 and *LIN28B* (chr6:104,950,183-104,964,925) RefSeq NM_001004317.4. Promoter denoted red; Proximal (Prox) enhancer sequences denoted orange; Distal (Dist) enhancer sequences denoted yellow. C) Schematic of genomic regulatory sequences identified from B). Fragments were cloned into the promoter-less *Renilla* luciferase construct – pGL4.79. The CMV-driven firefly luciferase construct pGL4.50 was used to normalize *Renilla.* Constructs tested in human ovarian teratocarcinoma cell line PA-1 (n=2).

Given the multifaceted spatiotemporal expression and function of LIN28A and LIN28B we sought to investigate the differential transcriptional regulation during key transition points throughout mammalian development. Publicly available chromatin immunoprecipitation sequencing (ChIP-seq) of the active histone marks in H1 human embryonic stem cells (hESC) (Ernst et al. 2011) (Fig.S1), armed us with the putative regulatory regions of *LIN28A* and *LIN28B.* We found several conserved genomic regulatory sequences for promoter regions (red), proximal (orange), and distal (yellow) enhancers of *LIN28B* maintained in human, chimp, mouse, frog and zebrafish, while the *LIN28A* promoter and enhancer regions preserved similarity in human, chimp, mouse and frog, but not zebrafish (Fig. 1B). Interestingly, only a single proximal enhancer region between exon 1 and exon 2 of *Lin28A* and *Lin28B* was conserved in lamprey and the other organisms. Next, we cloned these proximal fragments from H1 hESCs to determine the key regulatory sequences of *LIN28A* and *LIN28B*. Fragments were subcloned into luciferase constructs and then tested for activity in the embryonal carcinoma cell line, PA-1 (Fig. 1C). Not surprisingly, the -2kb fragment from the transcription start site, which contained both promoter and proximal enhancer sequences of both *Lin28A* and *Lin28B,* was more enzymatically active than the enhancer sequences alone. However, we did observe the +2/2.5kb fragment of *LIN28A,* which contained four proximal exonic enhancers, to also had high luciferase activity (Fig. 1C).

### Expression of *LIN28A* and *LIN28B* are differentially controlled by several master regulators of embryogenesis

Due to their capacity to maintain pluripotency states during early embryogenesis, the “Yamanaka factors” comprised of OCT4, KLF4, SOX2 and c-MYC (OSKM) are necessary and sufficient for induced pluripotent stem cell (iPSC) reprogramming (Okita et al. 2007). These transcription factors have been shown, together and individually, to regulate expression of *LIN28A* and *LIN28B* during stem/pluripotency maintenance and throughout lineage specification and differentiation. Sox2 regulates Lin28a, and to a lesser extent Lin28b, expression during neurogenesis in neural precursors (Cimadamore et al. 2013). The oncogene MYC is capable of transcriptionally upregulating LIN28B during cancer initiation, while LIN28A can replace MYC during iPSC generation (Yu et al. 2007; Chang et al. 2009). Finally, others found within the first 48 hours of OSKM expression during fibroblast reprogramming, the transcription factors cooperatively bind (Cacchiarelli et al. 2015; Soufi et al. 2015) and increase mRNA expression of *LIN28A,* and of *LIN28B*.

We investigated whether transcription factors known to control germ layer transitions to organogenesis could regulate the promoter/enhancer regions of human *LIN28A* and *LIN28B* (Fig. 2A). We conducted an *in vitro* high-throughput expression and luciferase reporter screen in HEK293 cells stably expressing the -2kb regulatory fragment driving *Renilla* luciferase (Fig. 2A-2C). As positive controls in our initial screen, we found the pluripotency factors OCT4, SOX2, KLF4, NANOG, MYC (Fig. 2D) were able to increase both the expression and luciferase activity of *LIN28A,* and to a somewhat lesser extent *LIN28B*, possibly due to basal *LIN28B* expression in HEK293 (Fig. 2B-2C). NANOG was able to increase luciferase activity, but not expression, of *LIN28B* (Fig. 2C). We found transcription factors important for stem/progenitor cell maintenance (such as MYCN and B-CATENIN), endoderm formation (such as FOXA2 and FOXA3), (Ang et al. 1993; Burtscher and Lickert 2009; Scheibner et al. 2021), mesoderm/neuromesodermal differentiation (via NOTCH and HOXB4) (Cooper et al. 2024), lung specification (such as NKX2-1) (Krude et al. 2002), and hematopoiesis (by MYB) (Soza-Ried et al. 2010) all controlled the key regulatory sequences of *LIN28A* and *LIN28B* (Fig. 2C). We focused on further validation of our screen using the pluripotent factors and those factors whose role during lung organogenesis were similarly crucial for neuronal differentiation. We also included SOX9, as its pleiotropic role in disease and organogenesis of several organs/tissues is well known (Chang et al. 2013; Rockich et al. 2013; Nikolic et al. 2017; Danopoulos et al. 2018; Balachandran and Narendran 2023; Aggarwal et al. 2024). Further targeted luciferase reporter experiments validated previous screening data and confirmed differential activation of the promoter/enhancer regions of human *LIN28A* and *LIN28B* by SOX9, NKX2-1, FOXA2, FOXA3, NOTCH (ICD), MYB, MYCN and B-CATENIN (Fig. 2E-2F and S2).

**Fig. 2.**
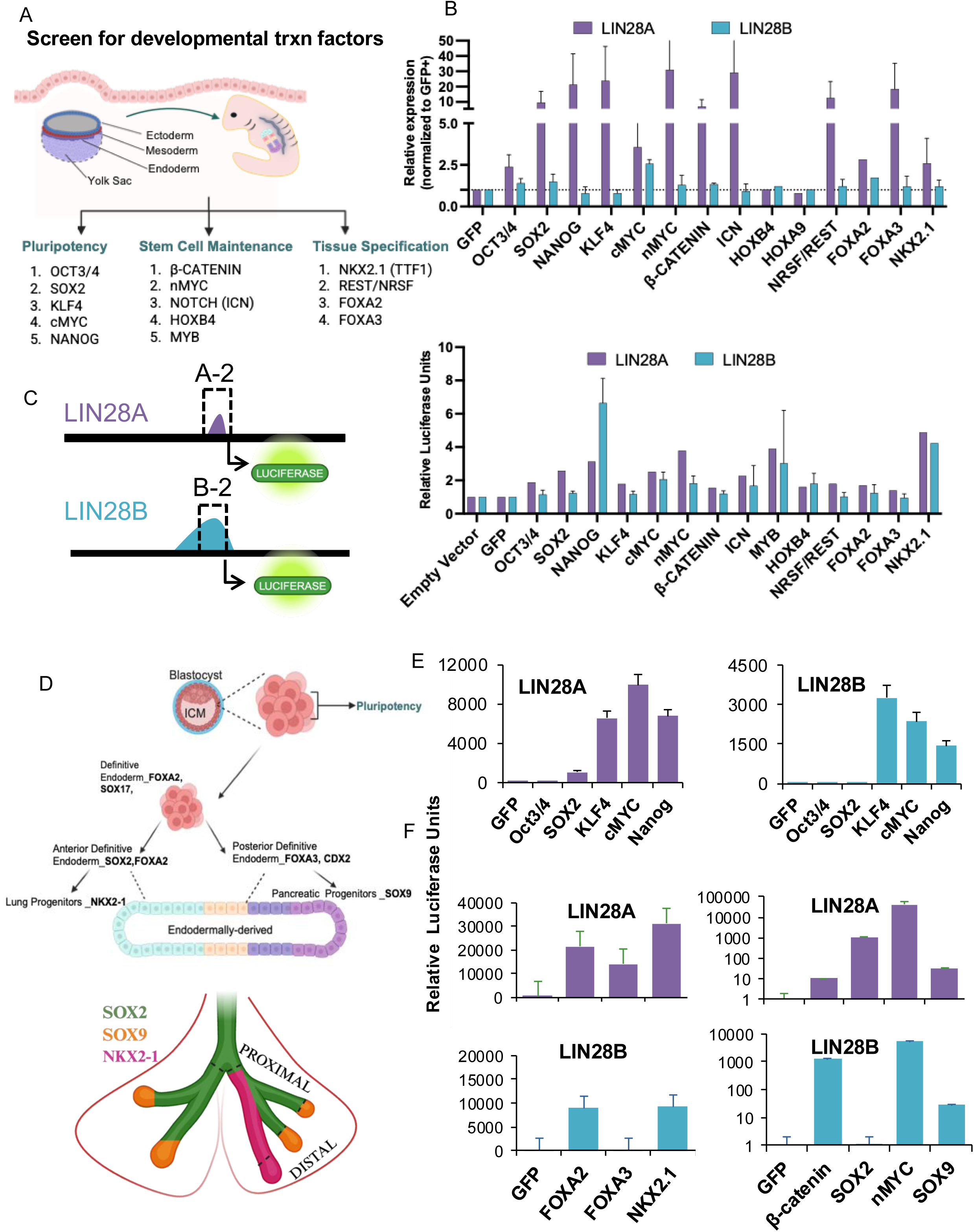
Transcriptional regulation of *LIN28A* and *LIN28B* by developmental transcription factors. A) Schematic of transcription factors that orchestrate germ layer formation and early embryonic lineage specification and differentiation. B) HEK293 were transiently transfected in 96-well format with the following factors: GFP (control), OCT3/4, SOX2, KLF4, NANOG, MYC, MYCN, SOX9, Intracellular cleaved NOTCH (ICN), B-CATENIN, HOXB4, HOXA9, NRSF/REST, FOXA2, FOXA3, and NKX2-1. RT-qPCR was conducted for human *LIN28A* and *LIN28B* (n=2 independent experiments in quadruplicate). C) Left: Schematic of -2kb fragments of *LIN28A* and *LIN28B* Right: HEK293 cells stably expressing CMV-driven firefly luciferase (selected with hygromycin) and -2kb fragments of *LIN28A* and *LIN28B* driving *Renilla* luciferase (selected with neomycin) were transiently transfected with transcription factors from B) that had 1.5 fold increase in expression (n=2 independent screening experiments, in triplicate, with the exception of NKX2-1, n=1 screen). D) Schematic of development of endoderm differentiation. Transient transfection of txrn factors into HEK293 with constructs stably expressing firefly and *Renilla* luciferase. Validation of a subset of factors that regulated human *LIN28A* and *LIN28B* promoter/enhancer sequences during screening experiments in panels B) and C) during E) pluripotency and F) endoderm/lung development.

Taken together, in addition to known regulation via OSKM, we uncovered several key developmental transcription factors known for multipotent stem/progenitor cell maintenance and germ layer formation that regulated *LIN28A* and *LIN28B* promoter/enhancer activity.

### Double knockout of Lin28a and Lin28b in developing lung epithelium mirror same mRNA targets in developing brain

We recently demonstrated that dKO of Lin28a/b in the lung epithelium have *let-7*-independent defects during early lung branching due to loss of mRNA-binding to *Etv5, Sox2,* and more importantly *Sox9.* (Osborne et al. 2021). Thus, our next steps were two-fold; first, we sought to determine in vivo, if there were feedback/feedforward regulation between Lin28 proteins and transcription factors from our initial screen. Our hypothesis, transcription factors upregulate Lin28a/b expression to enhance stabilization and translation of their own mRNAs to complete morphogenetic programs. Previous work from our group suggested Lin28a in the lung epithelium binds to and enhances the stability and translation of *Sox9* mRNA. Second, we investigated whether canonical Lin28-RBP motifs overlapped with the conserved 3’UTR *let-7-*miRNA binding motifs of early developmental transcription factors important for lung/endoderm and neuronal differentiation. Using a Sox2^eGFP^ reporter, we observed loss of Lin28a/b at the early onset of lung branching led to delayed lateral branching, which eventually led to decreased size and complexity of branches during later developmental time points (Fig.3A-3B). We recently reported decreased size of 50% of the dKO lungs at later time points compared to no difference in size at earlier times (Osborne et al. 2021), recapitulated here, suggesting possible proliferation defects (Fig.3A-3B). Previous studies have implicated LIN28A control of cell cycle progression via both *let-7*-dependent and independent mechanisms involving the regulation of cyclin and cyclin-dependent kinases (CDK) (Li et al. 2012). Evaluation of cyclin and CDK expression in E12.5 dKO lungs led to the conclusion that mRNA-mediated branching defects were not via these general cell cycle regulators (Fig.S3A-S3B). However, there was a significant decrease in gene expression of the proliferation marker, KI-67 at E12.5, which translated into decreased protein expression at E14.5 (Fig.3C-3D), suggesting proliferation defects could be partly responsible for size differences.

**Fig. 3.**
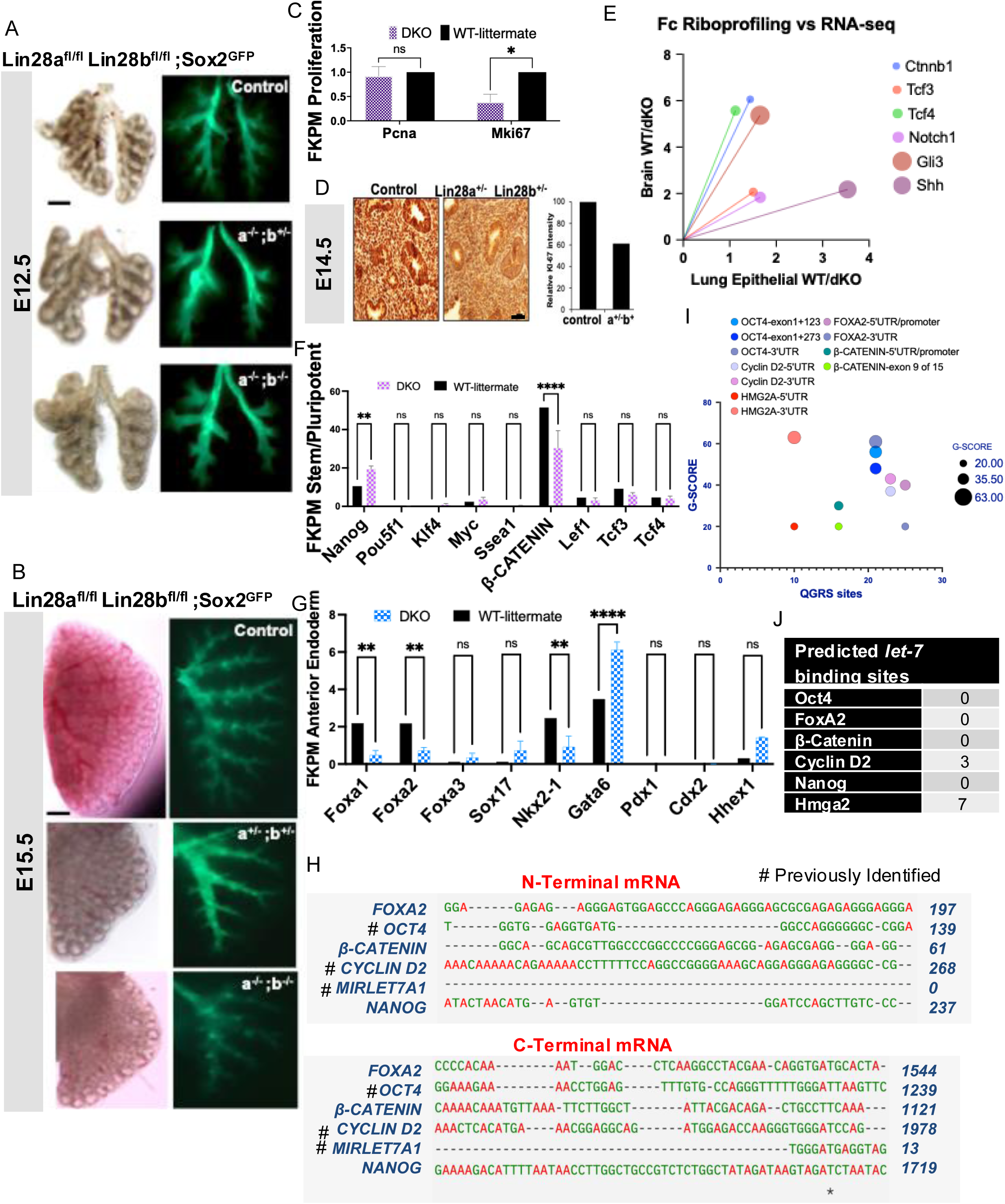
Loss of Lin28a/b in the developing lung leads to decreased mRNA expression of anterior endoderm/lung specification genes. Brightfield and fluorescent images of whole left lungs from wild type (WT) littermate or mutant (NKX2-1^creER^; Sox2^GFP^; Lin28a^+/-^;Lin28b^+/-^ (dHetKO), Lin28a^-/-^;Lin28b^+/-^ (aKO,bhet), or Lin28a^-/-^Lin28b^-/-^ (dKO) removed at A) E12.5 and B) E15.5 from dams given tamoxifen at E10.5. C) Bar graphs demonstrating Mki-67 gene expression from RNA-seq of the lung specific dKO of Lin28a/b (n=3) and WT. D) Left image: Immunohistochemistry staining of sectioned WT and dHetKO lungs for Ki-67. Right graph: quantification of Ki-67 intensity in WT and dHetKO. E) Fold change of mRNAs found during polyribosomal profiling in the Lin28a/b dKO/WT in brain (Herrlinger et al. 2019) compared to expression found in RNA-seq of the Lin28a/b dKO/WT in lung epithelium (Osborne et al. 2021). FKPM of genes involved in F) stem/pluripotency maintenance and G) endodermal differentiation from RNA-seq of the lung-specific Lin28a/b dKO/WT (n=3). H) Sequence alignment of select mRNAs from F) and G) performed in Clustal Omega. I) QGRS scores (G-Scores) compared to total number of QGRS of previously identified direct targets of Lin28a/b and genes of interest FOXA2 and B-CATENIN. J) Predicted *let-7* binding sites of known and putative Lin28-mRNA targets in TargetScan (Agarwal et al. 2015) https://www.targetscan.org/vert_80/. Data values are means of +/- S.D. Scale bars for: A) and B) are 0.5mm; for D) 50μm. *P* values for all graphs were determined using 2-way ANOVA, *P* values of significance: *≤ 0.05, **≤0.01, ***≤0.005, ****≤0.001.

The use and re-use of morphogenetic signaling pathways and their downstream gene regulatory networks coordinate the spatiotemporal development of all complex organs/tissues. The progenitors of each epithelial lung compartment in the mouse are marked by Sox2^+^ cells in the developing proximal lung and Sox9+ cells at the multipotent distal tips (Gontan et al. 2008; Que et al. 2009; Rawlins et al. 2009; Tompkins et al. 2009; Chang et al. 2013; Rockich et al. 2013; Wang et al. 2013; Alanis et al. 2014). Gene Ontology analysis from human embryonic distal (SOX9^+^/SOX2^+^) and proximal (SOX2^+^) lung progenitors interestingly indicate a neuronal signature enriched more in the SOX2^+^ proximal progenitors than the SOX9^+^/ SOX2^+^ distal progenitors (Nikolic et al. 2017). Given the known functions of the LIN28 proteins during neural development, neuronal differentiation, and recently during axonal regeneration in the adult mouse (Cimadamore et al. 2013; Yang et al. 2015; Wang et al. 2018; Nathan et al. 2020), we performed comparative analysis of mRNAs regulated by Lin28a/b in dKO lung and neuronal progenitors using our RNA-sequencing and previously published ribosomal profiling of the developing brain from identical developmental time points (Herrlinger et al. 2019). The mRNAs from signaling pathways including Notch, Wnt/B-Catenin, and Shh, together with their downstream transcriptional effectors were being actively translated in polyribosome fractions in the brain and differentially expressed in our lung samples (Fig.3E). RNA-sequencing of lung-specific dKO revealed expression levels of known Lin28-mRNA targets such as the stem/pluripotent factor Oct4 (*Pou5f1*) and the cell-cycle regulators Cyclins D2 (*Ccnd2*) and B1 (*Ccnb1*) did not change (Fig.3F and S3A and S3B). Surprisingly, we did observe derepression of *Nanog*, as well as repression of *B-Catenin* mRNAs in the lung-specific dKO (Fig.3F) that were not previously described as Lin28 or *let-7*-mRNA targets.

Next, we selected transcription factors responsible for lung/endoderm differentiation for which we observed regulation of the *LIN28A* and *LIN28B* enhancer/promoter regions. Genes critical for anterior foregut endoderm/lung specification such as *Foxa1, Foxa2,* and *Nkx2-1*, were significantly decreased expression upon dKO of Lin28a/b, while genes important for posterior foregut endoderm/hindgut formation such as *Foxa3, Pdx1, Cdx2* and *Hhex1* were unchanged (Fig.3G). Interestingly, expression of *Gata6*, shown to be important for endoderm fate specification of intestine, lung and pancreatic formation, was increased (Heslop et al. 2021) (Fig.3G). Multiple sequence alignments of known and potentially direct Lin28-mRNA targets revealed a curious observation. Two clusters formed, one at the 5’UTR N-terminus for those with relatively high guanine content and the other at the 3’UTR C-terminus with low to moderate guanine content (Fig.3H). Intriguingly, the 3’UTR mRNA cluster also grouped with motifs within the *let-7* miRNA whose binding by Lin28a/b CSD and the two CCHC zinc-finger domains result in inhibition and ultimate degradation (Viswanathan et al. 2008).

The ability of guanines to form 3D structures called G-quartets, which are composed of tetrad links stabilized via hydrogen bonds, has been associated with enhanced mRNA – stability, splicing, translation initiation and repression (Kikin et al. 2006). The bioinformatics tool, Quadruplex G-Rich Sequence (QGRS) mapper, was designed to predict these G-quartets 3D structures within mRNA sequences that are bound by RNA-binding proteins (Kikin et al. 2006). Utilizing QGRS mapper, we examined the presence of the canonical Lin28a RNA binding motif GGAGAU (conserved in both miRNA and mRNA), as well as G-rich quartet structures with the degenerate sequences AAGNNG, AAGNG(N), and (N)UGUG(N), where “N” could be any nucleotide (Wilbert et al. 2012; O’Day et al. 2015). High QGRS scores are sequences that are more likely to form a high number of stable G-quartets. Comparison of sequences from known LIN28-mRNA and -miRNA bound targets (*Mirlet7A1, Pou5f1,* and *Hmga2*) to those in our lung-specific dKO data set revealed moderate to high QGRS scores in *Foxa2* and *B-Catenin* (Fig.3I). Further analysis supported a differential in cluster formation whereby the 5’UTR/promoter regions of *Pou5f1, Foxa2,* and *B-Catenin* had higher QGRS scores and were not predicted targets of *let-7* miRNA (Fig.3H and 3J). However, *Ccnd2* and *Hmga2,* which are known targets of both Lin28 and *let-7,* had higher QGRS scores in the 3’UTR regions that aligned with the *let-7* miRNA motifs that are bound by Lin28a/b (Fig.3H and 3J). Taken together, these data suggest embryonic tissue and developing organs use not only transcriptionally controlled spatiotemporal means but also positional sequence-specific post-transcriptional mechanisms to fine-tune developmental timing.

Loss of Lin28 paralogs in neural progenitor cells (NPC) was previously shown to cause significant defects in neurulation, which included reduced rates of proliferation, failure of neural tube closure, microcephaly, and precocious differentiation of glial lineages (Shinoda et al. 2013b; Yang et al. 2015; Herrlinger et al. 2019). We used the neurosphere assay as our model of differentiation – where neural stem cells isolated from embryonic or adult brains form suspension spheroids, capable of limited self-renewal and multipotent differentiation (Fig.4A) (Gritti et al. 1996). We were able to repeat previously described defects in BDNF-induced neuronal differentiation at embryonic time points in vitro (Amen et al. 2017). Loss of Lin28a/b post-neurosphere formation (Fig. S4A) led to decreased complexity (measured by axonal projections and neuronal arborization) (Fig.4B), but not cell number (Fig. S4B and S4C). Furthermore, we observed dKO of Lin28a/b post-neurosphere formation did not prevent neuronal differentiation but did prevent extensive axonal arborization (Fig.4C). Genes previously identified to be downstream mRNA targets of Lin28a in NPCs include components of the Igf2-mTOR pathways–*Igf2, Igf2r1* and the Igf2 binding proteins (*Imps1,2,3*) and *Hmga2* (Yang et al. 2015). We examined the expression of these mRNAs in our lung-specific dKO of Lin28a/b. As we formerly observed, loss of Lin28a/b led to decreased expression of *Hmga2,* (Osborne et al. 2021), however, as we also previously observed, this was not dependent on *let-7* expression. Interestingly, we did observe a derepression of Igf2, but no significant changes in any of the other previously identified mRNAs in the pathway (Fig.4D).

**Fig. 4.**
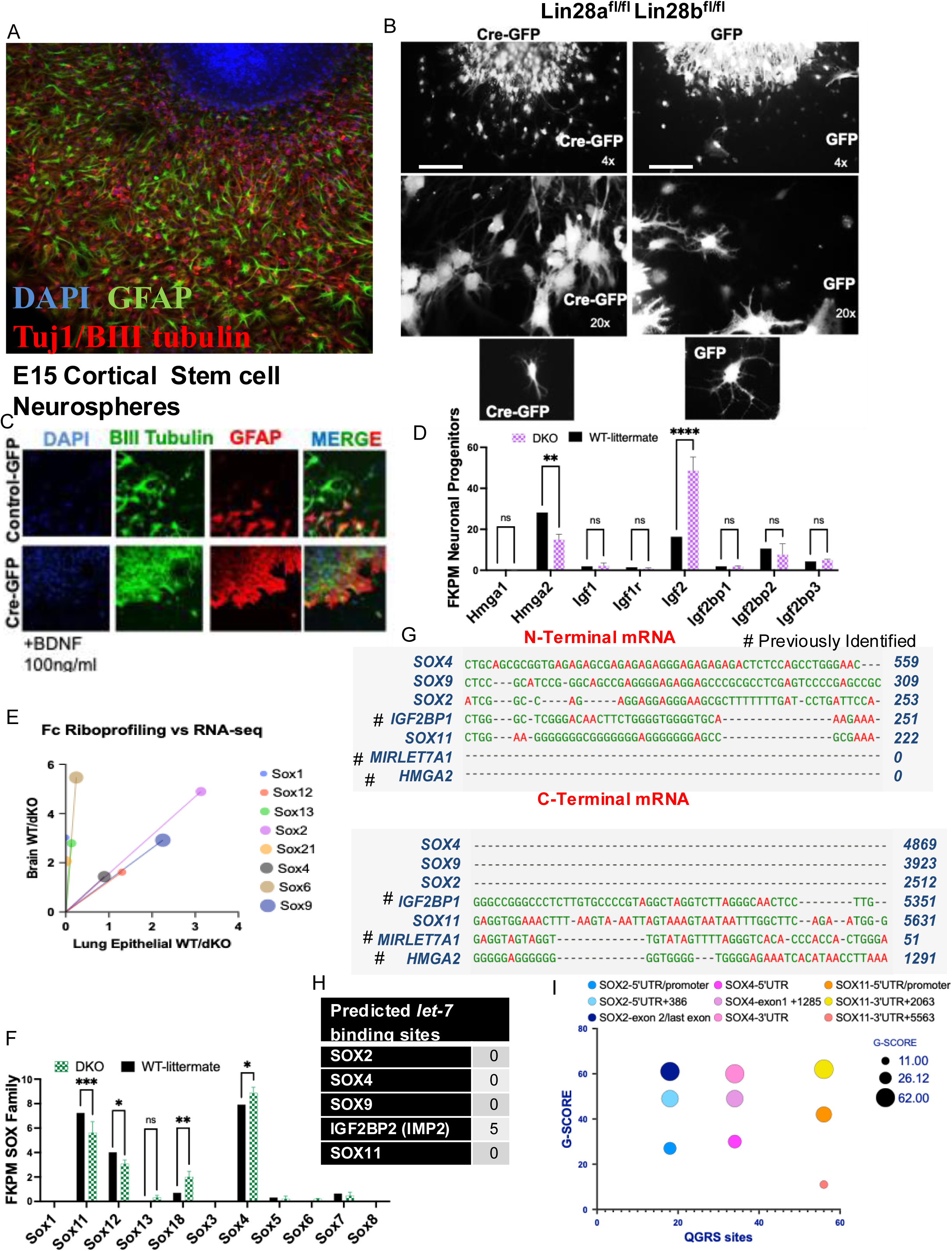
Loss of Lin28a/b in neural stem cells lead to decreased axonal projections and arborization. A) Neurospheres were derived from CD-1 E15 embryonic cortical neural stem cells. Spheroids were allowed to spontaneously differentiate in differentiation media (without BDNF or EGF). Progenitors of neurons and glial that migrated away from spheroids were then fixed and stained for Tuj1/BIII tubulin (neuronal) and GFAP (glia). Neurospheres were derived from Lin28a^floxed^Lin28b^floxed^ E12 brains. After sphere formation ex vivo, cultures were treated with Cre-GFP or CMV-GFP then directed to differentiate with the addition of B) 50ng/ml BDNF for 7 days or C)100ng/ml BDNF for 5 days then fixed and stained for Tuj1/BIII tubulin (neuronal) and GFAP (glia). D) FKPM of downstream mRNA targets of Lin28a/b in NPC from RNA-seq of the lung-specific Lin28a/b dKO/WT (n=3). E) Fold change of mRNAs found during polyribosome profiling in the Lin28a/b dKO/WT in brain (Herrlinger et al. 2019) compared to expression found in RNA-seq of the Lin28a/b dKO/WT in lung epithelium (Osborne et al. 2021). F) FKPM of Sox genes from RNA-seq of the lung-specific Lin28a/b dKO/WT (n=3). G) Sequence alignment of select Sox mRNAs performed in Clustal Omega. H) QGRS scores (G-Scores) compared to total number of QGRS of SOX2, SOX4, and SOX11 mRNAs. I) Predicted *let-7* binding sites of putative and known Lin28-mRNA targets in TargetScan (Agarwal et al. 2015) https://www.targetscan.org/vert_80/.

Given the cooperative feedback and feedforward regulation between the epithelial and mesenchymal pathways in the lung, we also investigated the expression of mesodermal lineage transcription factors, from our lung-specific dKO of Lin28a/b. In our whole lung lysates, we observed no change in mesodermal-promoting lineage transcription factors (Fig.S4D).

Sox proteins (specifically groups SoxB, SoxD, and SoxE) are well known for their regulatory functions during central and peripheral nervous system development (Kamachi and Kondoh 2013), in addition to the recently established roles of Sox2 and Sox9 in the developing lung epithelium (Rockich et al. 2013; Alanis et al. 2014). Since our previous data demonstrated Lin28-mRNA binding to *Sox2* and *Sox9* in the lung, we investigated the other Sox proteins, particularly those from the SoxC group (comprised of Sox4, Sox11, and Sox12), which are also shown to regulate neuronal differentiation (Mu et al. 2012). Brain-specific and lung-specific dKO of Lin28a/b showed several *Sox* mRNAs (including *Sox2, Sox9* and the SoxC group) being actively translated in polyribosome fractions in the brain and differentially expressed in the lung samples (Fig.4E). Subsequent analysis of the SoxC group in the lung-specific dKO of Lin28a/b demonstrated that *Sox11* and *Sox12* mRNA expression levels significantly decreased while *Sox4* increased (Fig.4F). Sequence alignments of *Sox2, Sox4, Sox9,* and *Sox11* mRNAs revealed 5’UTR N-terminal clusters with relatively moderate guanine content (Fig.4G). Only the *Sox11* 3’UTR C-terminus had low guanine content that aligned with the known mRNA targets of Lin28a/b protein and *let-7*-miRNA, including *Imps1/2* and *Hmga2,* though *Sox11* itself is not a predicted target of *let-7*-miRNA (Fig.4G and 4H). Finally, we found that unlike *Foxa2* and *B-Catenin* mRNAs, *Sox2, Sox4*, and *Sox11* (which are all intronless genes) had higher QGRS scores in the C-terminus/3’UTR regions, although none were predicted targets of the *let-7*-miRNA (Fig.4H and 4I). Altogether, while 3’UTR regions had higher QGRS scores, 5’UTR regions of *Sox2, Sox4*, and *Sox11* still had moderate scores. This suggests since the post-transcriptional regulation via Lin28-RBPs at the 3’UTR and 5’UTR regions are not in competition with 3’UTR *let-7-*miRNA binding sites, that there may be other secondary functions yet to be discovered that enables refinement of Lin28-mRNA binding and regulation (Fig.4I).

### *Lin28a* and *Lin28b* mRNA are involved in positive regulatory loops with B-catenin, Sox2 and Sox9 transcription factors *in vivo*

Previously, we observed that lung-specific dKO of Lin28a/b interrupted expression of *Etv5, Sox2* and *Sox9* and that Lin28a could bind to each of their mRNAs within the exonic regions directly (Osborne et al. 2021). Recently published data demonstrated that epithelial-specific knockout of B-Catenin led to decreased expression of Sox9 in distal tips, as well as branching defects (Ostrin et al. 2018). Considering we found B-CATENIN was able to: 1) differentially regulate the *LIN28A* and *LIN28B* enhancer/promoter regions (Fig.2F), and 2) dKO of Lin28a/b led to decreased *B-Catenin* mRNA expression (Fig.3F), we further investigated the presence of a feedforward regulatory loop between B-Catenin, Sox2, Sox9 and *Lin28a/b* mRNA in the lung epithelium in vivo. As we have described above, Sox2/SOX2 marks the embryonic mouse and human proximal stalks, while Sox9/SOX9 and SOX2 marks the distal tip regions, however B-Catenin, like Nkx2-1 marks stalk and tips (Mucenski et al. 2003; Cimadamore et al. 2013; Ostrin et al. 2018)(Fig. 5A). Wnt/B-Catenin signaling is required for lung/foregut progenitor specification– upstream of Fgf10 and its receptor Fgfr2– and necessary for proper distal-proximal airway formation, as loss leads to proximalization of the lung via ectopic Sox2 expression (Mucenski et al. 2003; Ostrin et al. 2018). Similar to previously described models of epithelial-specific knockout (Ostrin et al. 2018), we found loss of B-Catenin led to decreased expression of *Fgf10*, and increased expression of *Sox2* as well as no change in the downstream mediator of Shh signaling, *Gli1* (Fig. 5B). Interestingly, loss of B-Catenin led to decreased expression of *Lin28a* but not *Lin28b* (Fig. 5B). Together with Sox9, other transcription factors are also expressed at the distal tips, including MycN and Id2 (Okubo et al. 2005; Rawlins et al. 2009). While homozygous loss of Sox2 of genotypically pooled lungs led to modestly decreased expression of all genes examined, except *MycN,* epithelial-specific heterozygous and homozygous loss of Sox9 led to a dose-dependent decrease in the expression of *Lin28a* and *B-Catenin* but no considerable change in the levels of *Lin28b, Sox2, Bmp4,* or *MycN* (Fig.5B and 5C). Taken together, our data suggests the heterochronic programmers, Lin28a/b, control developmental transitions in early lung via enhanced mRNA stability and translation of key transcription factors that in turn transcriptionally upregulate *Lin28a,* and to a lesser extent *Lin28b,* expression (Fig.5E).

**Fig. 5.**
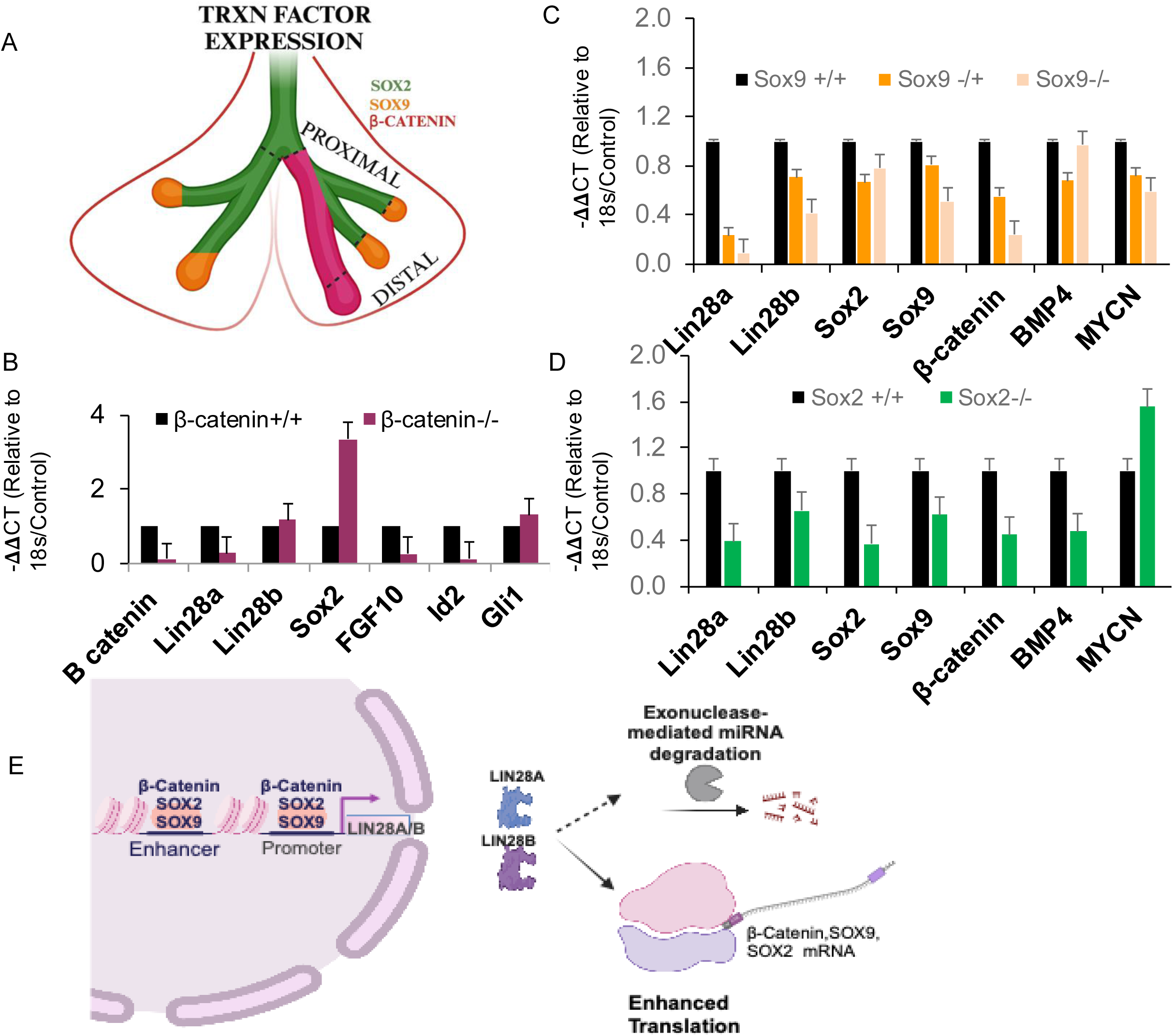
Transcription factors whose mRNA are regulated by Lin28 also regulate its expression in vivo. A) Schematic of the early developing lung, showing spatial expression of key transcription factors of the epithelium. Epithelial-specific knockout of B) *B-catenin*, C) *Sox9*, D) *Sox2* driven by *NKX2-1^creER^* administered with tamoxifen at E10.5. Lungs were isolated between E11.5 and E12.5. RNA was extracted, qRT-PCR was performed for the following genes: *Lin28a, Lin28b, Sox2, Sox9, Bmp4, B-CATENIN and Mycn, Fgf10, Id2* and *Gli1*. *Nkx2-1^creER^*; *B-catenin^fl/fl^*= 2 litters of pooled homozygous knockout & WT littermate lungs; *Sox9^fl/fl^*= 2 litters of pooled heterozygous, homozygous knockout & WT littermate lungs; *Sox2^fl/fl^*= 1 litter of pooled homozygous knockout & WT littermate lungs. E) Model of gene regulatory loops whereby transcriptional upregulation of Lin28a/b by transcription factors feeds into enhanced mRNA stability and/or maintenance of these same transcription factors.

## Discussion

The evolution of multicellularity necessitated the emergence of tissue-specific stem and progenitor cells. These specialized cells, via their specification and subsequent differentiation, facilitate the exponential growth and maturation of the organism. Heterochronic pathways, such as the lin-28/*let-7* axis, govern the timing and rate at which stem/progenitor cells stay in exponential growth phases, and when they exit to adapt different cell fates (Ambros 1997). We and others have shown that during early organogenesis in mammalian embryos, *let-7* miRNAs are not expressed until mid-gestation during the period when most stem/progenitor cells initiate differentiation (Yang et al. 2015; Yermalovich et al. 2019; Osborne et al. 2021). It has been suggested that during the early-to-mid gestational switch, Lin28 binds the stem-loop of the *let-7’s,* obstructing their processing and subsequent expression until progenitors are ready to differentiate. During the mid-to-late gestational switch at the initiation of differentiation, *let-7* in turn binds the 3′ UTRs of the *Lin28* mRNAs to suppress their translation, allowing differentiation to proceed (Piskounova et al. 2008; Rybak et al. 2008; Viswanathan et al. 2008). While there have been many investigations into the actions of the Lin28/*let-7* axis during mammalian development, few have fully investigated the early actions of the Lin28 paralogs prior to *let-7* expression (Moss et al. 1997). One study of note demonstrated the actions of the Lin28 paralogs in obesity/diabetes-resistance were due to both *let-7*-independent and -dependent mechanisms (Zhu et al. 2011). Zhu *et al*. demonstrated that several mRNAs in the insulin/PI3K-mTOR signaling pathway could have their translation enhanced by Lin28a/LIN28B and be suppressed by the *let-7* miRNAs. Our current study investigates the early transcriptional activation of Lin28a/b by the same factors we previously demonstrated to be direct mRNA targets.

In *C. elegans,* lin-28 loss of function led to precocious development and skipping the L2 stage completely (Moss et al. 1997). We previously determined double knockout of Lin28a and Lin28b in the lung epithelium leads to perinatal lethality, due to incomplete lung development (Osborne et al. 2021). Reminiscent of prior studies in *C. elegans*, we also found that Lin28 functions via two genetically distinct mechanisms during lung development, whereby during early embryogenesis, Lin28 binds mRNAs to enhance their translation, while throughout fetal maturation, binds to miRNA to facilitate their degradation before the onset of differentiation (Balzer et al. 2010; Vadla et al. 2012; Osborne et al. 2021). Considering that the expression of Lin28a occurs in a wave during lung development, but not during the development of other organs or tissues examined, leaves room for speculation that there may be interesting biology yet to be uncovered during late-gestational stages of lung alveologenesis (Shinoda et al. 2013b; Tu et al. 2015; Yang et al. 2015; Yermalovich et al. 2019; Osborne et al. 2021).

Previously, we demonstrated that Lin28a could bind exonic regions of the mRNAs *Sox2, Sox9, Etv5, Gli1,* and *Bmp4,* in the lung, heart, brain, kidney, and liver (Osborne et al. 2021). The canonical Lin28a and Lin28b RNA binding motif GGAGAU is found to be conserved in both miRNA and mRNA, where the CSD preferentially binds “UGAU”, and the two CCHC zinc-finger domains have higher affinity for “GGAG” (Ustianenko et al. 2018; Tan et al. 2021). Recent studies indicate that the mass of RNAs bound by LIN28 in mammalian ESCs and human cancer cell lines are protein-coding mRNAs, not miRNAs (Tan et al. 2021). However, while many human cancer cell lines have high expression of either LIN28A or LIN28B, many also have moderate to high levels of the *let-7* miRNA family (Powers et al. 2016). This complicated paradox is absent throughout early organogenesis, where we and others have demonstrated *let-7* miRNAs are not fully expressed until mid-late gestation, at the onset of differentiation (Balzer et al. 2010; Vadla et al. 2012; Cimadamore et al. 2013; Yang et al. 2015; Yermalovich et al. 2019; Osborne et al. 2021). Here we report that in addition to spatiotemporal gene regulatory loops between LIN28 and downstream RNAs, in mRNAs specifically there are 5’UTR and 3’UTR position-dependent post-transcriptional mechanisms that are used to fine tune developmental timing. A better understanding of the distinct mechanisms of how G-quartet 3D structures form at 5’UTRs compared to 3’UTRs in vivo could be harnessed to improve precision medicine during adult regeneration, particularly given the recently described actions of Lin28 in axonal regeneration (Wang et al. 2018).

Crosstalk and feedback/feedforward manipulation of morphogenetic pathways are critical for the proper development of all metazoans. Pleiotropic transcriptional effectors such as Sox9, are critical for development and adult stem cell maintenance of numerous organs and tissues, and are activated by several pathways such as Fgf9/Fgf10-Fgfr2b in the lung, pancreas, and testes; Pdgfrα in the nervous system; Wnt in the intestine and pancreas, Notch1 in the testes and retina, as well as Shh in the skin (Chang et al. 2013; Rockich et al. 2013; Danopoulos et al. 2018). Loss of Sox9 in the lung epithelium leads to severe branching defects and incomplete differentiation of the alveolar lineages (Chang et al. 2013; Rockich et al. 2013; Danopoulos et al. 2018). We found that Lin28a and Sox9 formed a *let-7*-independent feedback loop during lung branching by demonstrating that heterozygous loss of *Sox9* could partially rescue certain phenotypes observed with overexpression of *Lin28a* (Osborne et al. 2021). We report here that in addition to the capability of LIN28 protein to bind and enhance the stability of *SOX9* mRNA (Osborne et al. 2021); SOX2, SOX9 and B-CATENIN proteins can also transactivate the *cis-* regulatory elements of the human *LIN28A* and *LIN28B* genes and control expression of mouse *Lin28a* in the lung epithelium.

## Material and Methods

### Mouse Models

All mouse experiments were conducted following the Institutional Animal Care and Use Committee at Boston Children’s Hospital and UT Southwestern Medical Center that is accredited by AAALAC. Generation of the conditional double knockout (Hibar et al.) was as previously described(Zhu et al. 2011; Shinoda et al. 2013b; Osborne et al. 2021). The *NKX2.1-*creER (014552), *Lox-stop-Lox-tdTomato* (007914), and *Sox9* conditional KO mice (013106), *Sox2* conditional KO (013093), *Ctnnb1* conditional KO mice (004152) *Sox2*-GFP (017592) were purchased from The Jackson Laboratory. The mice used in the study were embryonic ages during E11.5-E18.5. Tamoxifen (Sigma Aldrich, 579000) was diluted in corn oil to 20mg/ml and administered intraperitoneally at a dose of 0.068mg/g per mouse weight in grams and at E10.5-11. Wildtype littermate controls (without expression of Cre recombinase) were used for all experiments n=pooled embryos from two different dams for *Sox9, Sox2, Ctnnb1* conditional homozygous KO and one for *Sox9* conditional heterozygous KO.

### Cell culture

Embryonal ovarian PA-1 teratocarcinoma cells (CRL-1572) and HEK293 cells (CRL-1573) purchased from ATCC were maintained in DMEM+10%FBS at 37 degrees and 5%CO2. Cells were transfected with all plasmid constructs using Lipofectamine 2000 (Invitrogen, 11668019) and lysed using Passive Lysis buffer (Promega, E1941) for luciferase assay or TRIzol^TM^ Reagent (Invitrogen, 15596026) for downstream mRNA analysis.

### qRT-PCR, RNA-sequencing and analysis

RNAs were isolated from lungs and cell lines using TRIzol^TM^ Reagent (Invitrogen, 15596026). For mRNA analysis, cDNA was prepared from 2 μg RNA using miScript II RT Kit (Qiagen, 218160) for high throughput analysis and 5μg RNA using iScript^TM^ Advanced cDNA Synthesis Kit (Bio-Rad, 1725037). For qRT-PCR, primer oligos for Lin28a (forward-AGC TTG CAT TCC TTG GCA TGA TGG; reverse-AGG CGG TGG AGT TCA CCT TTA AGA) and Lin28b (forward-TTT GGC TGA GGA GGT AGA CTG CAT; reverse-ATG GAT CAG ATG TGG ACT GTG CGA) were synthesized by IDT. Primers for mRNAs: *Shh, Sox2, Sox9, Bmp4, Smad1, β catenin*, and *MycN* were from BioRad (PrimePCR SYBR), primers for *FGF10, ETV5,* and *Gli1* were as previously described in (Herriges et al. 2015). The raw and analyzed RNA-seq data in detailed method is available through Gene Expression Omnibus (GEO) with the ID GSE93571 as detailed in our previous manuscript (Osborne et al. 2021). Ribosomal profiling seq data set_ GEO under accession number GSE131536. From manuscript (Herrlinger et al. 2019).

### Reporter Assays

Human LIN28A and LIN28B regulatory elements (including promoter, distal and proximal enhancers) were cloned from H1 human embryonic stem cell genomic DNA (Wi-Cell). Regulatory elements were defined by using UCSC genome browser ID: hg38 (Perez et al. 2024) and ChIP-seq deposited on UCSC genome browser of active histone marks, H3K27ac, H3K4me1, and H3K4me3 (Ernst et al. 2011). LIN28A/LIN28B elements were cloned: -3kb to +2.5kb for A; and -3kb to +2kb from the transcriptional start site (Lango Allen et al.). Promoter and enhancer fragments of LIN28A/B (A-2kb, A+2 kb, A+2.5kb; B-2 kb, B-13kb and B+2kb) were PCR subcloned into the 5’KPNI-3’XHOI sites of pGL4.79-neomycin-Renilla luciferase, while the pGL4.50 CMV-driven-hygromycin-Firefly luciferase was used as control. Stable HEK293 cell pools expressing pGL4.49/pGL4.50 HEK293, selected in neomycin and hygromycin respectively, were used to screen and validate transcription factors (Table S1) that activated Renilla luciferase and increased mRNA expression of LIN28A and Lin28B. Dual luciferase activity was monitored using the Dual-Luciferase^®^ Reporter Assay System (Promega, E1910), and results were normalized to GFP and empty vector expressing control samples. High-throughput reporter screen in HEK293 cells was performed on the BioTek Synergy HTX Multimode Reader (Agilent) using dual reagent injector kinetic assay mode. Screen analysis was done using Gen5 software (Agilent). Validation of luciferase reporter activity in regulatory fragments in PA-1 and HEK293 cells was performed on the Centro LB 963 microplate luminometer (Berthold Technologies GmbH & Co.KG) and analysis was performed in Microsoft Excel and GraphPad Prism 10.

### Neurosphere Assays

Embryonic Brains from E12-E14.5 Lin28a*^floxed^*Lin28b*^floxed^* pups were dissected and dissociated in DNase Ι (0.5mg/ml) then re-suspended in HBSS/L15 media (1:1). For generation of neurospheres, cells were plated at a density of 10,000-100,000 cell/well in Costar^®^ 6-well Clear Flat Bottom Ultra-Low Attachment Multiple Well plate (Corning, 3471). All cells were re-suspended in DMEM/F-12: Neurobasal^TM^ Medium (5:3 mix; Gibco, 21103049), bFGF (20ng/ml; R&D systems, 3139-FB), N-2 Supplement (1%; Gibco, 17502048), B-27 Supplement (2%; Gibco, 17504044), β-Mercaptoethanol (Gibco, 21985023), 50 µM final. Self-renewal media (same as initiation) with addition of: EGF 20 ng/ml (R&D systems, 2028-EG), chicken egg extract 10%. Differentiation media: DMEM/F12: Neurobasal^TM^ Medium (5:3 mix; GIBCO), FBS 5%, bFGF (10ng/ml), N-2 Supplement (1%), B-27 Supplement (2%), β-Mercaptoethanol, 50 µM final with or without BDNF 50-100ng/ml (Santa Cruz Biotechnology, sc-4554). Neurospheres grown for 2-3 days in differentiation media were then dissociated and additionally grown in BDNF 100ng/ml for 5-7 days. Lin28a/b were knocked out post-neurosphere formation via adenoviral infection with Ad-Cre-GFP adenovirus (Vector Biolabs, 1700) or Ad-GFP adenovirus (Vector Biolabs, 1060).

### Sequence Alignment and QGRS Mapper

RNA Sequence alignments and analysis of guanine (G)-rich (G-rich) regions predicted to be LIN28-mRNA/miRNA binding sites was performed using the FASTA/NCBI of human mRNA sequences below: NC_000009.12:94175957-94176036 Homo sapiens chromosome 9, GRCh38.p14 Primary Assembly (MIRLET7A1); NM_002701.6 Homo sapiens POU class 5 homeobox 1 (POU5F1), transcript variant 1, mRNA (OCT4); NM_024865.4 Homo sapiens Nanog homeobox (NANOG), transcript variant 1, mRNA; NM_003106.4 Homo sapiens SRY-box transcription factor 2 (SOX2), mRNA; NM_003483.6 Homo sapiens high mobility group AT-hook 2 (HMGA2), transcript variant 1, mRNA; NM_021784.5 Homo sapiens forkhead box A2 (FOXA2), transcript variant 1, mRNA; NM_001904.4 Homo sapiens catenin beta 1 (CTNNB1), transcript variant 1, mRNA (Beta-CATENIN); Z46629.1 Homo sapiens SOX9 mRNA; NM_003107.3 Homo sapiens SRY-box transcription factor 4 (SOX4), mRNA; NM_003108.4 Homo sapiens SRY-box transcription factor 11 (SOX11). Multiple sequence alignments were done using Clustal Omega: https://www.ebi.ac.uk/jdispatcher/msa/clustalo. Identification of novel compared to known G-rich motifs was done using the bioinformatics tool: http://bioinformatics.ramapo.edu/QGRS/ to obtain G-scores. The G-rich motifs were identified using identification of G-tetrads in quadruplexes and then given a score based on the number of stable guanine-quadruplexes potentially formed.

### Tissue preparation, immunostaining, and antibodies

Embryos were removed from timed pregnant mice that were anesthetized via the open-drop method of isoflurane exposure followed by cervical dislocation and/or ketamine/xylazine. Lungs were dissected and fixed in 10% formalin overnight. For sectioned immunohistochemical staining, fixed lungs were moved to 70% ethanol. The protocol for immunohistochemical staining was performed as previously published (Tu et al. 2015). For immunofluorescent staining, fixed cells were incubated in blocking serum (PBS with 5% normal donkey serum (Sigma Aldrich, D9663), and 0.5% Triton X-100 (Sigma Aldrich) for 1 hour at room temperature and then incubated with primary antibodies diluted in blocking serum at 4℃ overnight. The following day, the cells were washed with PBS three times at room temperature and then incubated with secondary antibodies and DAPI diluted in blocking serum for 1 hour at room temperature. Antibodies and reagents: KI-67 (clone SP6), catalog no. ab16667, lot GR155005-1, Abcam; DAPI catalog no. D1306, Invitrogen; GFAP, catalog no. ab5804, lot 2388830, Millipore; BIII Tubulin/Tuj1, (clone 2G10) catalog no. MA1-118, Invitrogen.

## Acknowledgements

We would like to thank and acknowledge George Q. Daley for mentorship and financial support via NIH R01GM107536. We would also like to thank, Patricia M. Sousa and Alena V. Yermalovich for assistance with maintenance of mouse colony and Areum Han for initial RNA-seq analysis of previous manuscript (Osborne et al. 2021); members of the Daley lab and Osborne lab, and Michael Reese for constructive reading of the manuscript. J.K.O. was supported as a post-doc by Burroughs Wellcome Fund, as a P.I. by the V Foundation V2021-028; CPRIT (Cancer Prevention Research Institute of Texas) start-up grant number, RR210016. S.F.L. was supported by Cancer Prevention and Research Institute of Texas (CPRIT) Pre-doctoral Training Grant number, RP210041. B.K. was partially funded by CPRIT Post-doctoral Training Grant number, RP210041.

## Author Contributions

J.K.O wrote the manuscript; M.F.O and S.F.L edited manuscript. J.K.O, I.T., M.F.O, E.S. S.B., C.G., B.K., R.G.R Performed experiments. E.L.R., help with bioinformatics analysis.

---

**Supplemental Fig. 1.**
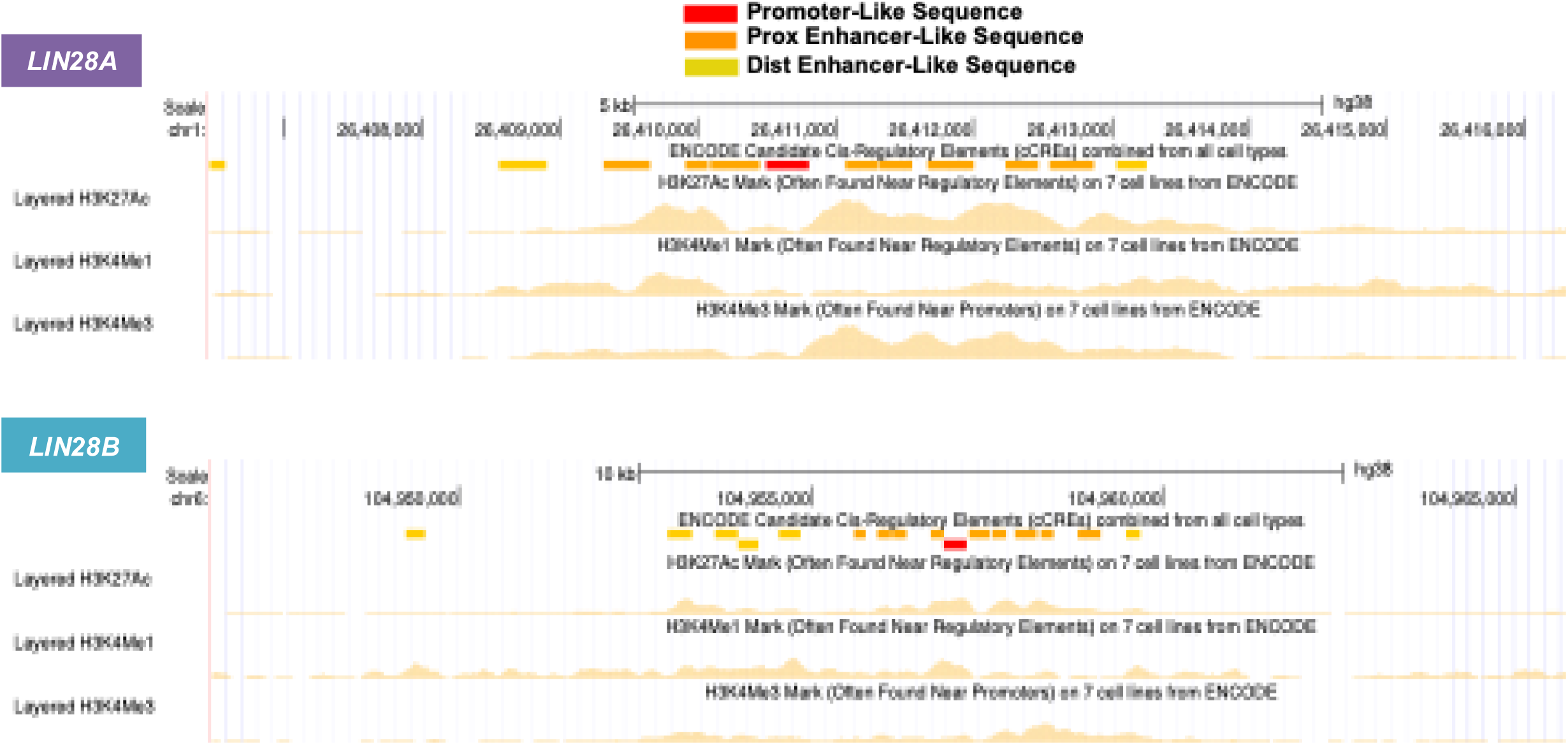
UCSC genome browser-defined epigenetic marks of the *LIN28A* and *LIN28B* regulatory regions. Active histone marks, H3K27ac, H3K4me1, and H3K4me3 in H1 human embryonic stem cells (hESC)(Ernst et al. 2011) of putative regulatory sequences, determined via chromatin immunoprecipitation sequencing (ChIP-seq) in the UCSC genome browser (Perez et al. 2024). Assembly used was hg38 for *LIN28A* (chr1:26,406,455-26,417,607) RefSeq NM_024674.6 and *LIN28B* (chr6:104,950,183-104,964,925) RefSeq NM_001004317.4. Promoter denoted red; Proximal (Prox) enhancer sequences denoted orange; Distal (Dist) enhancer sequences denoted yellow.

**Supplemental Fig. 2.**
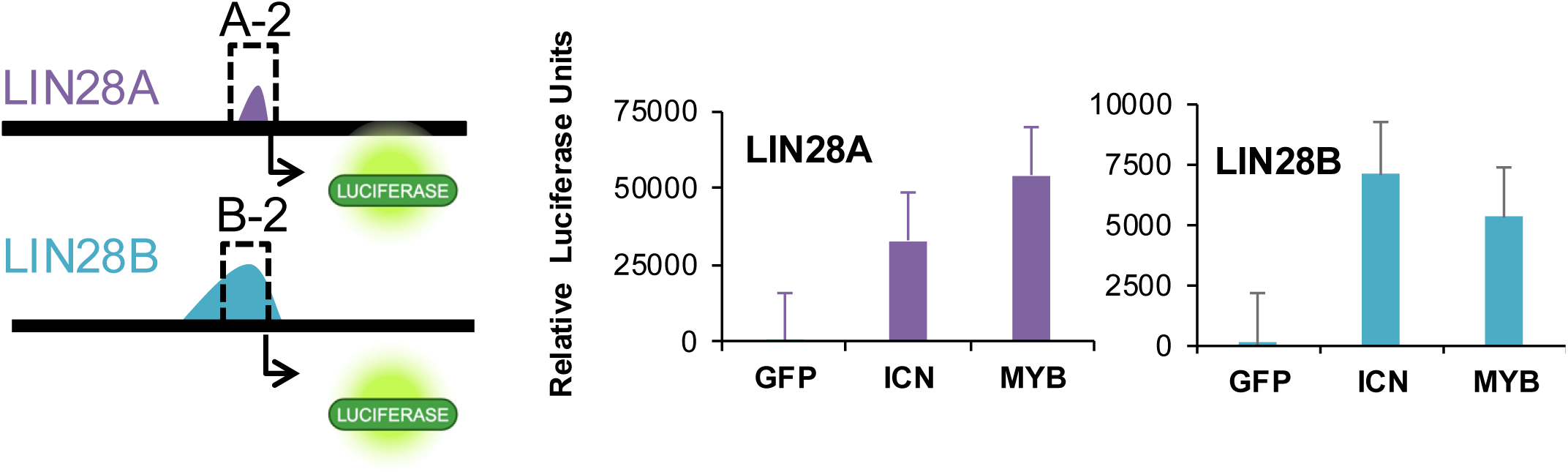
**Transcriptional regulation of *LIN28A* and *LIN28B*** Transcriptional regulation of *LIN28A* and *LIN28B* Validation of a subset of factors that regulated human *LIN28A* and *LIN28B* promoter/enhancer sequences during screening experiments in Figure 2B, for MYB and the intracellular cleaved domain of NOTCH1.

**Supplemental Fig. 3.**
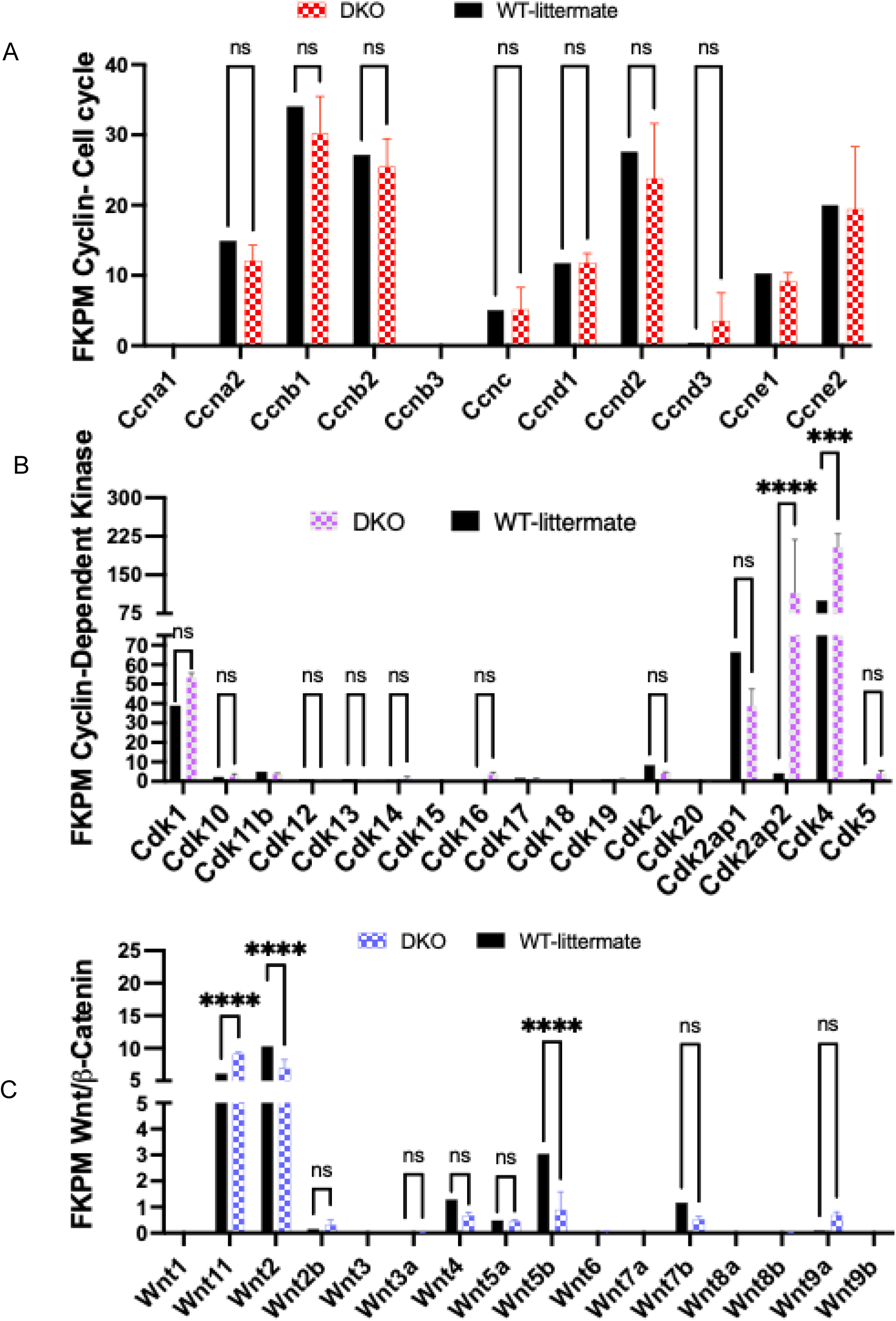
FKPM gene expression values of Cyclins, Cyclin-dependent Kinases, and Wnt family members. Expression of genes found in RNA-seq of the Lin28a/b dKO/WT in lung epithelium A) *Cyclin* genes B)*Cyclin-dependent Kinases* C) Family of *Wnt* ligands (Osborne et al. 2021).

**Supplemental Fig. 4.**
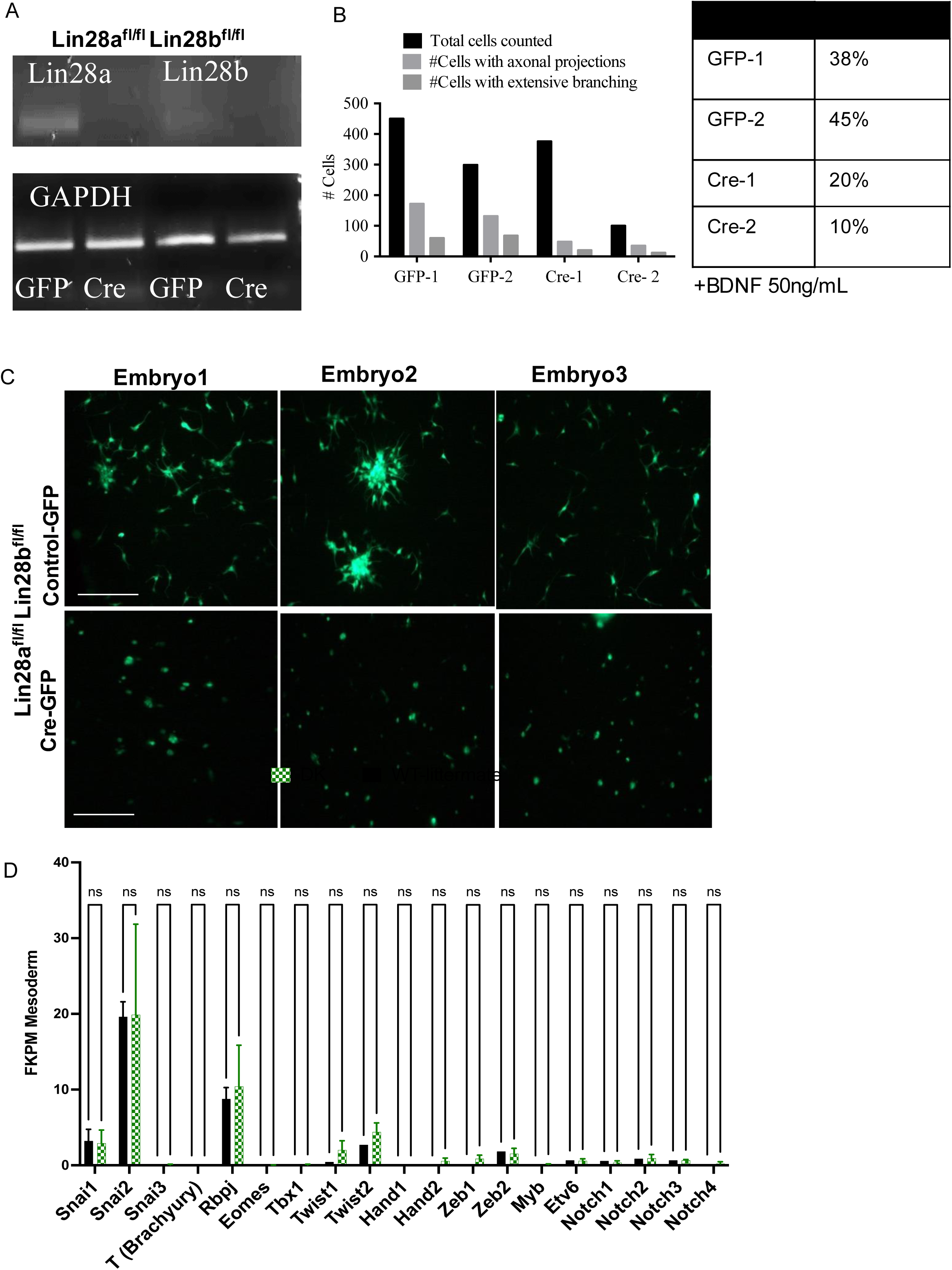
Characterization of the Lin28 double knockout in neural stem cells. Neurospheres derived from Lin28a^floxed^Lin28b^floxed^ E12 whole brains expressing CMV-GFP control adenovirus or Cre-Recombinase adenovirus: A) agarose gel of RT-PCR B) quantification of C C) BDNF-treated mechanically dissociated spheroids after 7 days in culture on TC-treated 96-well glass bottom plates.

**Table S1.**
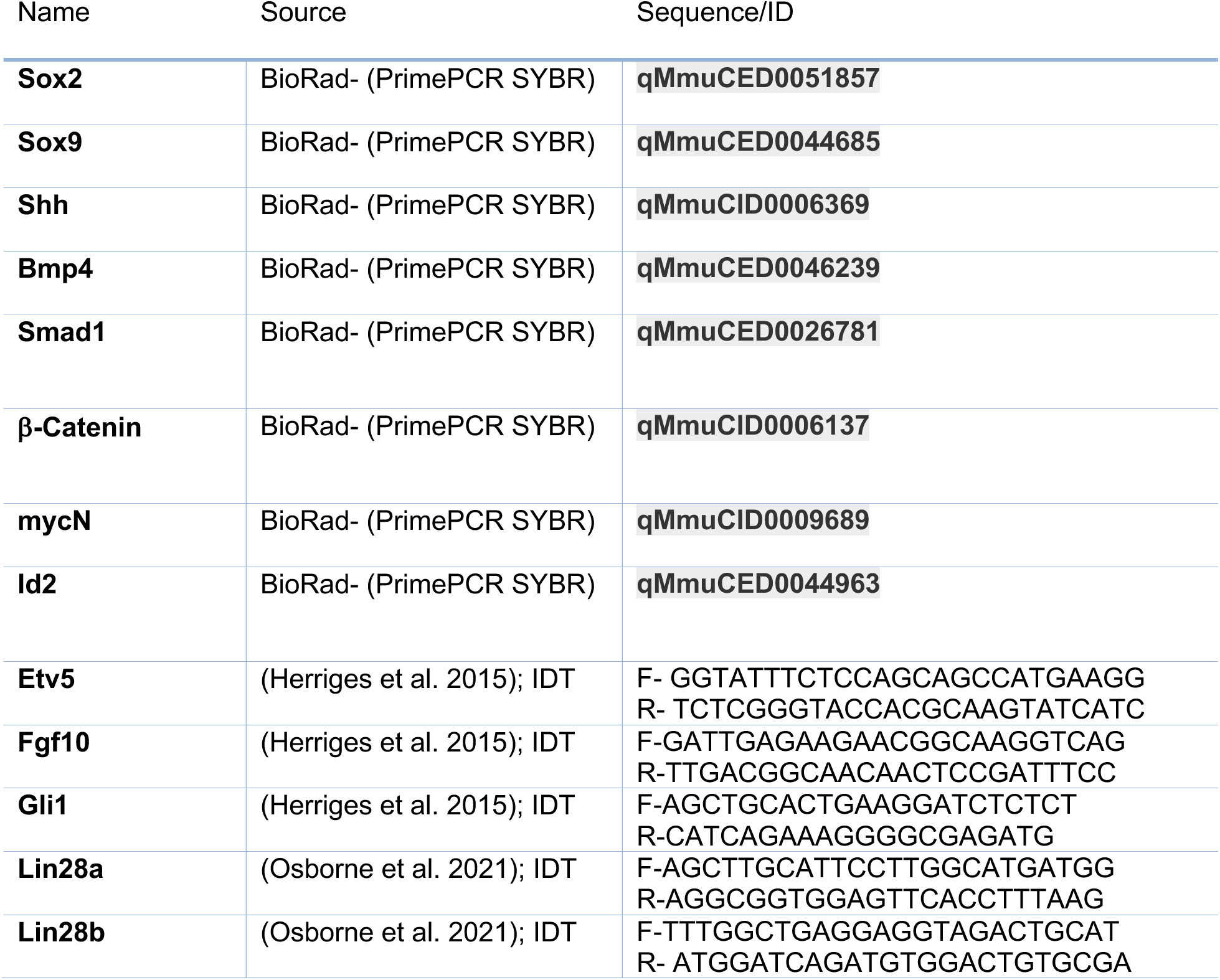
Primer Data.

**Table S2.**
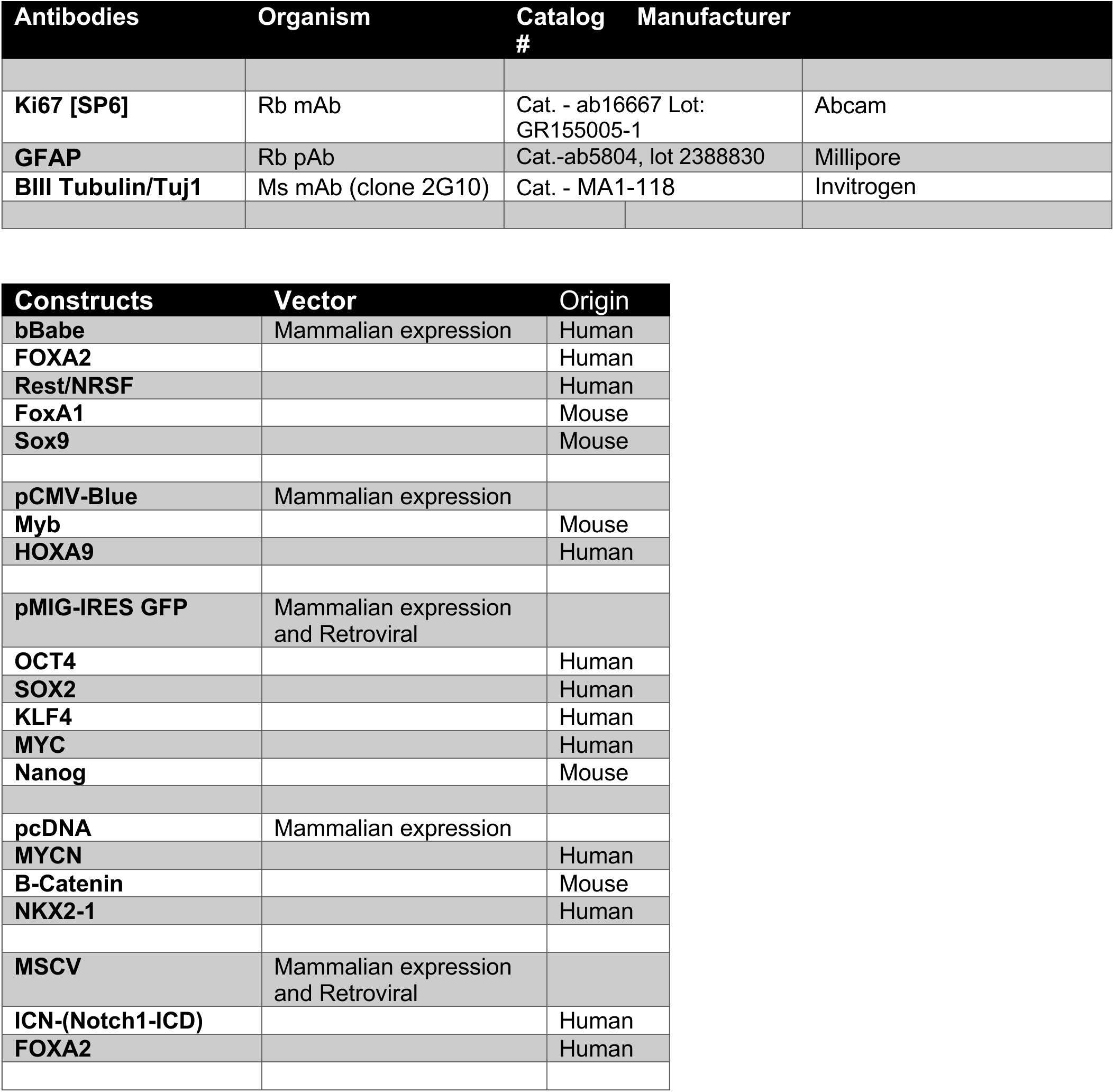
Constructs and Antibodies.

## References

Agarwal V, Bell GW, Nam JW, Bartel DP. 2015. Predicting effective microRNA target sites in mammalian mRNAs. Elife 4.

Aggarwal S, Wang Z, Rincon Fernandez Pacheco D, Rinaldi A, Rajewski A, Callemeyn J, Van Loon E, Lamarthee B, Covarrubias AE, Hou J et al. 2024. SOX9 switch links regeneration to fibrosis at the single-cell level in mammalian kidneys. Science 383: eadd6371.

Alanis DM, Chang DR, Akiyama H, Krasnow MA, Chen J. 2014. Two nested developmental waves demarcate a compartment boundary in the mouse lung. Nat Commun 5: 3923.

Ambros V. 1997. Heterochronic Genes. in C elegans II (eds. nd, DL Riddle, T Blumenthal, BJ Meyer, JR Priess), Cold Spring Harbor (NY).

Amen AM, Ruiz-Garzon CR, Shi J, Subramanian M, Pham DL, Meffert MK. 2017. A Rapid Induction Mechanism for Lin28a in Trophic Responses. Mol Cell 65: 490–503 e497.

Ang SL, Wierda A, Wong D, Stevens KA, Cascio S, Rossant J, Zaret KS. 1993. The formation and maintenance of the definitive endoderm lineage in the mouse: involvement of HNF3/forkhead proteins. Development 119: 1301–1315.

Balachandran S, Narendran A. 2023. The Developmental Origins of Cancer: A Review of the Genes Expressed in Embryonic Cells with Implications for Tumorigenesis. Genes (Basel) 14.

Balzer E, Heine C, Jiang Q, Lee VM, Moss EG. 2010. LIN28 alters cell fate succession and acts independently of the let-7 microRNA during neurogliogenesis in vitro. Development 137: 891–900.

Burtscher I, Lickert H. 2009. Foxa2 regulates polarity and epithelialization in the endoderm germ layer of the mouse embryo. Development 136: 1029–1038.

Cacchiarelli D, Trapnell C, Ziller MJ, Soumillon M, Cesana M, Karnik R, Donaghey J, Smith ZD, Ratanasirintrawoot S, Zhang X et al. 2015. Integrative Analyses of Human Reprogramming Reveal Dynamic Nature of Induced Pluripotency. Cell 162: 412–424.

Cai WY, Wei TZ, Luo QC, Wu QW, Liu QF, Yang M, Ye GD, Wu JF, Chen YY, Sun GB et al. 2013. The Wnt-beta-catenin pathway represses let-7 microRNA expression through transactivation of Lin28 to augment breast cancer stem cell expansion. J Cell Sci 126: 2877–2889.

Chang DR, Martinez Alanis D, Miller RK, Ji H, Akiyama H, McCrea PD, Chen J. 2013. Lung epithelial branching program antagonizes alveolar differentiation. Proc Natl Acad Sci U S A 110: 18042–18051.

Chang TC, Zeitels LR, Hwang HW, Chivukula RR, Wentzel EA, Dews M, Jung J, Gao P, Dang CV, Beer MA et al. 2009. Lin-28B transactivation is necessary for Myc-mediated let-7 repression and proliferation. Proc Natl Acad Sci U S A 106: 3384–3389.

Cimadamore F, Amador-Arjona A, Chen C, Huang CT, Terskikh AV. 2013. SOX2-LIN28/let-7 pathway regulates proliferation and neurogenesis in neural precursors. Proc Natl Acad Sci U S A 110: E3017–3026.

Cooper F, Souilhol C, Haston S, Gray S, Boswell K, Gogolou A, Frith TJR, Stavish D, James BM, Bose D et al. 2024. Notch signalling influences cell fate decisions and HOX gene induction in axial progenitors. Development 151.

Danopoulos S, Alonso I, Thornton ME, Grubbs BH, Bellusci S, Warburton D, Al Alam D. 2018. Human lung branching morphogenesis is orchestrated by the spatiotemporal distribution of ACTA2, SOX2, and SOX9. Am J Physiol Lung Cell Mol Physiol 314: L144–L149.

Ernst J, Kheradpour P, Mikkelsen TS, Shoresh N, Ward LD, Epstein CB, Zhang X, Wang L, Issner R, Coyne M et al. 2011. Mapping and analysis of chromatin state dynamics in nine human cell types. Nature 473: 43–49.

Faas L, Warrander FC, Maguire R, Ramsbottom SA, Quinn D, Genever P, Isaacs HV. 2013. Lin28 proteins are required for germ layer specification in Xenopus. Development 140: 976–986.

Gontan C, de Munck A, Vermeij M, Grosveld F, Tibboel D, Rottier R. 2008. Sox2 is important for two crucial processes in lung development: branching morphogenesis and epithelial cell differentiation. Dev Biol 317: 296–309.

Gritti A, Parati EA, Cova L, Frolichsthal P, Galli R, Wanke E, Faravelli L, Morassutti DJ, Roisen F, Nickel DD et al. 1996. Multipotential stem cells from the adult mouse brain proliferate and self-renew in response to basic fibroblast growth factor. J Neurosci 16: 1091–1100.

Gundermann DG, Martinez J, De Kervor G, Gonzalez-Pinto K, Larrain J, Faunes F. 2019. Overexpression of Lin28a delays Xenopus metamorphosis and down-regulates albumin independently of its translational regulation domain. Dev Dyn 248: 969–978.

Guo Y, Chen Y, Ito H, Watanabe A, Ge X, Kodama T, Aburatani H. 2006. Identification and characterization of lin-28 homolog B (LIN28B) in human hepatocellular carcinoma. Gene 384: 51–61.

Herriges JC, Verheyden JM, Zhang Z, Sui P, Zhang Y, Anderson MJ, Swing DA, Zhang Y, Lewandoski M, Sun X. 2015. FGF-Regulated ETV Transcription Factors Control FGF-SHH Feedback Loop in Lung Branching. Dev Cell 35: 322–332.

Herriges M, Morrisey EE. 2014. Lung development: orchestrating the generation and regeneration of a complex organ. Development 141: 502–513.

Herrlinger S, Shao Q, Yang M, Chang Q, Liu Y, Pan X, Yin H, Xie LW, Chen JF. 2019. Lin28-mediated temporal promotion of protein synthesis is crucial for neural progenitor cell maintenance and brain development in mice. Development 146.

Heslop JA, Pournasr B, Liu JT, Duncan SA. 2021. GATA6 defines endoderm fate by controlling chromatin accessibility during differentiation of human-induced pluripotent stem cells. Cell Rep 35: 109145.

Hibar DP, Stein JL, Renteria ME, Arias-Vasquez A, Desrivieres S, Jahanshad N, Toro R, Wittfeld K, Abramovic L, Andersson M, et al. 2015. Common genetic variants influence human subcortical brain structures. Nature 520: 224–229.

Kamachi Y, Kondoh H. 2013. Sox proteins: regulators of cell fate specification and differentiation. Development 140: 4129–4144.

Kikin O, D’Antonio L, Bagga PS. 2006. QGRS Mapper: a web-based server for predicting G-quadruplexes in nucleotide sequences. Nucleic Acids Res 34: W676–682.

Kimura S, Hara Y, Pineau T, Fernandez-Salguero P, Fox CH, Ward JM, Gonzalez FJ. 1996. The T/ebp null mouse: thyroid-specific enhancer-binding protein is essential for the organogenesis of the thyroid, lung, ventral forebrain, and pituitary. Genes Dev 10: 60–69.

Krude H, Schutz B, Biebermann H, von Moers A, Schnabel D, Neitzel H, Tonnies H, Weise D, Lafferty A, Schwarz S et al. 2002. Choreoathetosis, hypothyroidism, and pulmonary alterations due to human NKX2-1 haploinsufficiency. J Clin Invest 109: 475–480.

Lango Allen H, Estrada K, Lettre G, Berndt SI, Weedon MN, Rivadeneira F, Willer CJ, Jackson AU, Vedantam S, Raychaudhuri S et al. 2010. Hundreds of variants clustered in genomic loci and biological pathways affect human height. Nature 467: 832–838.

Li C, Sako Y, Imai A, Nishiyama T, Thompson K, Kubo M, Hiwatashi Y, Kabeya Y, Karlson D, Wu SH et al. 2017. A Lin28 homologue reprograms differentiated cells to stem cells in the moss Physcomitrella patens. Nat Commun 8: 14242.

Li N, Zhong X, Lin X, Guo J, Zou L, Tanyi JL, Shao Z, Liang S, Wang LP, Hwang WT et al. 2012. Lin-28 homologue A (LIN28A) promotes cell cycle progression via regulation of cyclin-dependent kinase 2 (CDK2), cyclin D1 (CCND1), and cell division cycle 25 homolog A (CDC25A) expression in cancer. J Biol Chem 287: 17386–17397.

Minoo P, Su G, Drum H, Bringas P, Kimura S. 1999. Defects in tracheoesophageal and lung morphogenesis in Nkx2.1(-/-) mouse embryos. Dev Biol 209: 60–71.

Monaghan AP, Kaestner KH, Grau E, Schutz G. 1993. Postimplantation expression patterns indicate a role for the mouse forkhead/HNF-3 alpha, beta and gamma genes in determination of the definitive endoderm, chordamesoderm and neuroectoderm. Development 119: 567–578.

Moss EG, Lee RC, Ambros V. 1997. The cold shock domain protein LIN-28 controls developmental timing in C. elegans and is regulated by the lin-4 RNA. Cell 88: 637–646.

Moss EG, Tang L. 2003. Conservation of the heterochronic regulator Lin-28, its developmental expression and microRNA complementary sites. Dev Biol 258: 432–442.

Mu L, Berti L, Masserdotti G, Covic M, Michaelidis TM, Doberauer K, Merz K, Rehfeld F, Haslinger A, Wegner M et al. 2012. SoxC transcription factors are required for neuronal differentiation in adult hippocampal neurogenesis. J Neurosci 32: 3067–3080.

Mucenski ML, Wert SE, Nation JM, Loudy DE, Huelsken J, Birchmeier W, Morrisey EE, Whitsett JA. 2003. beta-Catenin is required for specification of proximal/distal cell fate during lung morphogenesis. J Biol Chem 278: 40231–40238.

Nathan FM, Ohtake Y, Wang S, Jiang X, Sami A, Guo H, Zhou FQ, Li S. 2020. Upregulating Lin28a Promotes Axon Regeneration in Adult Mice with Optic Nerve and Spinal Cord Injury. Mol Ther 28: 1902–1917.

Nikolic MZ, Caritg O, Jeng Q, Johnson JA, Sun D, Howell KJ, Brady JL, Laresgoiti U, Allen G, Butler R et al. 2017. Human embryonic lung epithelial tips are multipotent progenitors that can be expanded in vitro as long-term self-renewing organoids. Elife 6.

O’Day E, Le MTN, Imai S, Tan SM, Kirchner R, Arthanari H, Hofmann O, Wagner G, Lieberman J. 2015. An RNA-binding Protein, Lin28, Recognizes and Remodels G-quartets in the MicroRNAs (miRNAs) and mRNAs It Regulates. J Biol Chem 290: 17909–17922.

Okita K, Ichisaka T, Yamanaka S. 2007. Generation of germline-competent induced pluripotent stem cells. Nature 448: 313–317.

Okubo T, Knoepfler PS, Eisenman RN, Hogan BL. 2005. Nmyc plays an essential role during lung development as a dosage-sensitive regulator of progenitor cell proliferation and differentiation. Development 132: 1363–1374.

Osborne JK, Kinney MA, Han A, Akinnola KE, Yermalovich AV, Vo LT, Pearson DS, Sousa PM, Ratanasirintrawoot S, Tsanov KM et al. 2021. Lin28 paralogs regulate lung branching morphogenesis. Cell Rep 36: 109408.

Ostrin EJ, Little DR, Gerner-Mauro KN, Sumner EA, Rios-Corzo R, Ambrosio E, Holt SE, Forcioli-Conti N, Akiyama H, Hanash SM et al. 2018. beta-Catenin maintains lung epithelial progenitors after lung specification. Development 145.

Ouchi Y, Yamamoto J, Iwamoto T. 2014. The heterochronic genes lin-28a and lin-28b play an essential and evolutionarily conserved role in early zebrafish development. PLoS One 9: e88086.

Perez G, Barber GP, Benet-Pages A, Casper J, Clawson H, Diekhans M, Fischer C, Gonzalez JN, Hinrichs AS, Lee CM et al. 2024. The UCSC Genome Browser database: 2025 update. Nucleic Acids Res.

Piskounova E, Viswanathan SR, Janas M, LaPierre RJ, Daley GQ, Sliz P, Gregory RI. 2008. Determinants of microRNA processing inhibition by the developmentally regulated RNA-binding protein Lin28. J Biol Chem 283: 21310–21314.

Powers JT, Tsanov KM, Pearson DS, Roels F, Spina CS, Ebright R, Seligson M, de Soysa Y, Cahan P, Theissen J et al. 2016. Multiple mechanisms disrupt the let-7 microRNA family in neuroblastoma. Nature 535: 246–251.

Que J, Luo X, Schwartz RJ, Hogan BL. 2009. Multiple roles for Sox2 in the developing and adult mouse trachea. Development 136: 1899–1907.

Rawlins EL, Clark CP, Xue Y, Hogan BL. 2009. The Id2+ distal tip lung epithelium contains individual multipotent embryonic progenitor cells. Development 136: 3741–3745.

Rockich BE, Hrycaj SM, Shih HP, Nagy MS, Ferguson MA, Kopp JL, Sander M, Wellik DM, Spence JR. 2013. Sox9 plays multiple roles in the lung epithelium during branching morphogenesis. Proc Natl Acad Sci U S A 110: E4456–4464.

Rybak A, Fuchs H, Smirnova L, Brandt C, Pohl EE, Nitsch R, Wulczyn FG. 2008. A feedback loop comprising lin-28 and let-7 controls pre-let-7 maturation during neural stem-cell commitment. Nat Cell Biol 10: 987–993.

Scheibner K, Schirge S, Burtscher I, Buttner M, Sterr M, Yang D, Bottcher A, Ansarullah, Irmler M, Beckers J et al. 2021. Epithelial cell plasticity drives endoderm formation during gastrulation. Nat Cell Biol 23: 692–703.

Serls AE, Doherty S, Parvatiyar P, Wells JM, Deutsch GH. 2005. Different thresholds of fibroblast growth factors pattern the ventral foregut into liver and lung. Development 132: 35–47.

Shinoda G, De Soysa TY, Seligson MT, Yabuuchi A, Fujiwara Y, Huang PY, Hagan JP, Gregory RI, Moss EG, Daley GQ. 2013a. Lin28a regulates germ cell pool size and fertility. Stem Cells 31: 1001–1009.

Shinoda G, Shyh-Chang N, Soysa TY, Zhu H, Seligson MT, Shah SP, Abo-Sido N, Yabuuchi A, Hagan JP, Gregory RI et al. 2013b. Fetal deficiency of lin28 programs life-long aberrations in growth and glucose metabolism. Stem Cells 31: 1563–1573.

Soufi A, Garcia MF, Jaroszewicz A, Osman N, Pellegrini M, Zaret KS. 2015. Pioneer transcription factors target partial DNA motifs on nucleosomes to initiate reprogramming. Cell 161: 555–568.

Soza-Ried C, Hess I, Netuschil N, Schorpp M, Boehm T. 2010. Essential role of c-myb in definitive hematopoiesis is evolutionarily conserved. Proc Natl Acad Sci U S A 107: 17304–17308.

Sun Z, Yu H, Zhao J, Tan T, Pan H, Zhu Y, Chen L, Zhang C, Zhang L, Lei A et al. 2022. LIN28 coordinately promotes nucleolar/ribosomal functions and represses the 2C-like transcriptional program in pluripotent stem cells. Protein Cell 13: 490–512.

Tan FE, Sathe S, Wheeler EC, Yeo GW. 2021. Non-microRNA binding competitively inhibits LIN28 regulation. Cell Rep 36: 109517.

Tompkins DH, Besnard V, Lange AW, Wert SE, Keiser AR, Smith AN, Lang R, Whitsett JA. 2009. Sox2 is required for maintenance and differentiation of bronchiolar Clara, ciliated, and goblet cells. PLoS One 4: e8248.

Tu HC, Schwitalla S, Qian Z, LaPier GS, Yermalovich A, Ku YC, Chen SC, Viswanathan SR, Zhu H, Nishihara R et al. 2015. LIN28 cooperates with WNT signaling to drive invasive intestinal and colorectal adenocarcinoma in mice and humans. Genes Dev 29: 1074–1086.

Ustianenko D, Chiu HS, Treiber T, Weyn-Vanhentenryck SM, Treiber N, Meister G, Sumazin P, Zhang C. 2018. LIN28 Selectively Modulates a Subclass of Let-7 MicroRNAs. Mol Cell 71: 271–283 e275.

Vadla B, Kemper K, Alaimo J, Heine C, Moss EG. 2012. lin-28 controls the succession of cell fate choices via two distinct activities. PLoS Genet 8: e1002588.

Viswanathan SR, Daley GQ, Gregory RI. 2008. Selective blockade of microRNA processing by Lin28. Science 320: 97–100.

Vogt EJ, Meglicki M, Hartung KI, Borsuk E, Behr R. 2012. Importance of the pluripotency factor LIN28 in the mammalian nucleolus during early embryonic development. Development 139: 4514–4523.

Wang XW, Li Q, Liu CM, Hall PA, Jiang JJ, Katchis CD, Kang S, Dong BC, Li S, Zhou FQ. 2018. Lin28 Signaling Supports Mammalian PNS and CNS Axon Regeneration. Cell Rep 24: 2540–2552 e2546.

Wang Y, Tian Y, Morley MP, Lu MM, Demayo FJ, Olson EN, Morrisey EE. 2013. Development and regeneration of Sox2+ endoderm progenitors are regulated by a Hdac1/2-Bmp4/Rb1 regulatory pathway. Developmental cell 24: 345–358.

Wells JM, Melton DA. 1999. Vertebrate endoderm development. Annu Rev Cell Dev Biol 15: 393–410.

West JA, Viswanathan SR, Yabuuchi A, Cunniff K, Takeuchi A, Park IH, Sero JE, Zhu H, Perez-Atayde A, Frazier AL et al. 2009. A role for Lin28 in primordial germ-cell development and germ-cell malignancy. Nature 460: 909–913.

Wilbert ML, Huelga SC, Kapeli K, Stark TJ, Liang TY, Chen SX, Yan BY, Nathanson JL, Hutt KR, Lovci MT et al. 2012. LIN28 binds messenger RNAs at GGAGA motifs and regulates splicing factor abundance. Mol Cell 48: 195–206.

Yang M, Yang SL, Herrlinger S, Liang C, Dzieciatkowska M, Hansen KC, Desai R, Nagy A, Niswander L, Moss EG et al. 2015. Lin28 promotes the proliferative capacity of neural progenitor cells in brain development. Development 142: 1616–1627.

Yermalovich AV, Osborne JK, Sousa P, Han A, Kinney MA, Chen MJ, Robinton DA, Montie H, Pearson DS, Wilson SB et al. 2019. Lin28 and let-7 regulate the timing of cessation of murine nephrogenesis. Nat Commun 10: 168.

Yu J, Vodyanik MA, Smuga-Otto K, Antosiewicz-Bourget J, Frane JL, Tian S, Nie J, Jonsdottir GA, Ruotti V, Stewart R et al. 2007. Induced pluripotent stem cell lines derived from human somatic cells. Science 318: 1917–1920.

Zhang J, Ratanasirintrawoot S, Chandrasekaran S, Wu Z, Ficarro SB, Yu C, Ross CA, Cacchiarelli D, Xia Q, Seligson M et al. 2016. LIN28 Regulates Stem Cell Metabolism and Conversion to Primed Pluripotency. Cell Stem Cell 19: 66–80.

Zhu H, Shyh-Chang N, Segre AV, Shinoda G, Shah SP, Einhorn WS, Takeuchi A, Engreitz JM, Hagan JP, Kharas MG et al. 2011. The Lin28/let-7 axis regulates glucose metabolism. Cell 147: 81–94.

